# NHE6-Depletion Corrects ApoE4-Mediated Synaptic Impairments and Reduces Amyloid Plaque Load

**DOI:** 10.1101/2021.03.22.436385

**Authors:** Theresa Pohlkamp, Xunde Xian, Connie H Wong, Murat Durakoglugil, Gordon C Werthmann, Takaomi Saido, Bret M Evers, Charles L White, Jade Connor, Robert E Hammer, Joachim Herz

## Abstract

Apolipoprotein E4 (ApoE4) is the most important and prevalent risk factor for late-onset Alzheimer’s disease (AD). The isoelectric point of ApoE4 matches the pH of the early endosome (EE), causing its delayed dissociation from ApoE receptors and hence impaired endolysosomal trafficking, disruption of synaptic homeostasis and reduced amyloid clearance. We have shown that enhancing endosomal acidification by inhibiting the EE-specific sodium-hydrogen exchanger NHE6 restores vesicular trafficking and normalizes synaptic homeostasis. Remarkably and unexpectedly, loss of NHE6 effectively suppressed amyloid deposition even in the absence of ApoE4, suggesting that accelerated acidification of early endosomes caused by the absence of NHE6 occludes the effect of ApoE on amyloid plaque formation. NHE6 suppression or inhibition may thus be a universal, ApoE-independent approach to prevent amyloid buildup in the brain. These findings suggest a novel therapeutic approach for the prevention of AD by which partial NHE6 inhibition reverses the ApoE4 induced endolysosomal trafficking defect and reduces amyloid.

## INTRODUCTION

ApoE is the principal lipid transport protein in the brain. Three different ApoE isoforms are common in humans: ApoE2 (ɛ2), ApoE3 (ɛ3), and ApoE4 (ɛ4). Each ApoE4 allele reduces the age of Alzheimer’s disease (AD) onset by approximately three to five years compared to ApoE3 homozygotes, which comprise ∼80% of the human population (Roses, 1994; Sando et al., 2008). By contrast and by comparison, ApoE2 is protective against AD (Corder et al., 1994; Panza et al., 2000; West et al., 1994). ApoE is an arginine-rich protein and a major component of very-low density lipoproteins (Shore and Shore, 1973). The number of positively charged arginine residues differs between the three human isoforms due to two single nucleotide polymorphisms in the ApoE gene. The most common isoform, ApoE3, has a charge neutral cysteine at amino acid position 112 and an arginine at position 158. The second most common isoform, ApoE4, has two arginines, while the less frequent ApoE2 has two cysteines at these respective positions. The positively charged arginines raise the net charge and thus the isoelectric point (IEP) of the protein. The IEP of ApoE2 is the lowest (5.9), ApoE3 has an intermediate IEP of 6.1 and the IEP of ApoE4 is ∼6.4 (Ordovas et al., 1987). For cargo delivery, ApoE binds to lipoprotein receptors and undergoes endocytosis and recycling. Endocytic subcompartments become progressively more acidic, and the pH of these compartments is regulated by the opposing functions of vesicular ATP-dependent proton pumps (vATPase) and proton leakage channels (Na^+^/H^+^ exchangers, NHEs). Early endocytic vesicles are slightly acidic (pH, ∼6.4), which facilitates ligand/receptor dissociation. Lysosomes are highly acidic (pH 4 to 5), which is required for the digestion of endocytosed biomolecules (**Figure 1A**) (Casey et al., 2010; Naslavsky and Caplan, 2018). For maturation of the early endosomes (EE) and entry into the next sorting stage, ligand/receptor dissociation is required. The pH-dependent release of ApoE from its receptor in the EE is important for endosomal maturation and cargo delivery (Yamamoto et al., 2008) for the ability of endosomal content to rapidly recycle to the cell surface (Heeren et al., 1999; Nixon, 2017). The early endosomal pH, which triggers ligand-receptor dissociation, closely matches the IEP of ApoE4 (Xian et al., 2018). Loss of net surface charge at the IEP is accompanied by reduced solubility in an aqueous environment, leading to impaired dissociation of ApoE4 from its receptors (Xian et al., 2018) and aided by a greater propensity of ApoE4 to form a molten globule configuration under acidic conditions (Morrow et al., 2002). Dysregulation of endolysosomal trafficking by ApoE4 causes an age dependent increase in EE number and size (Nuriel et al., 2017). Based on these observations, we have proposed a model in which destabilization of ApoE4 in the acidic EE environment, combined with a greater propensity for self-association, results in delayed detachment from its receptors (**Figure 1B**). Subsequent endosomal swelling through K^+^, Na^+^ and H_2_O influx further impairs cargo delivery, receptor recycling, and ligand re-secretion. Importantly, in neurons, ApoE and its receptor Apoer2 travel together with glutamate receptors through the endosomal recycling pathway (Chen et al., 2010; Xian et al., 2018). Rapid endocytosis and subsequent recycling of synaptic receptors is triggered by the synaptic homeostatic modulator and Apoer2 ligand Reelin (Hiesberger et al., 1999; Trommsdorff et al., 1999). We previously showed that ApoE4, in contrast to ApoE3 and ApoE2, prevents Reelin-mediated glutamate receptor insertion at the synapse, a state we refer to as ApoE4-mediated Reelin resistance (Chen et al., 2010; Durakoglugil et al., 2009; Lane-Donovan and Herz, 2017; Lane-Donovan et al., 2014; Xian et al., 2018). Reduction of endosomal pH and increasing the differential to the ApoE4 IEP abolishes this effect *in vitro* (Xian et al., 2018).

**Figure 1:**
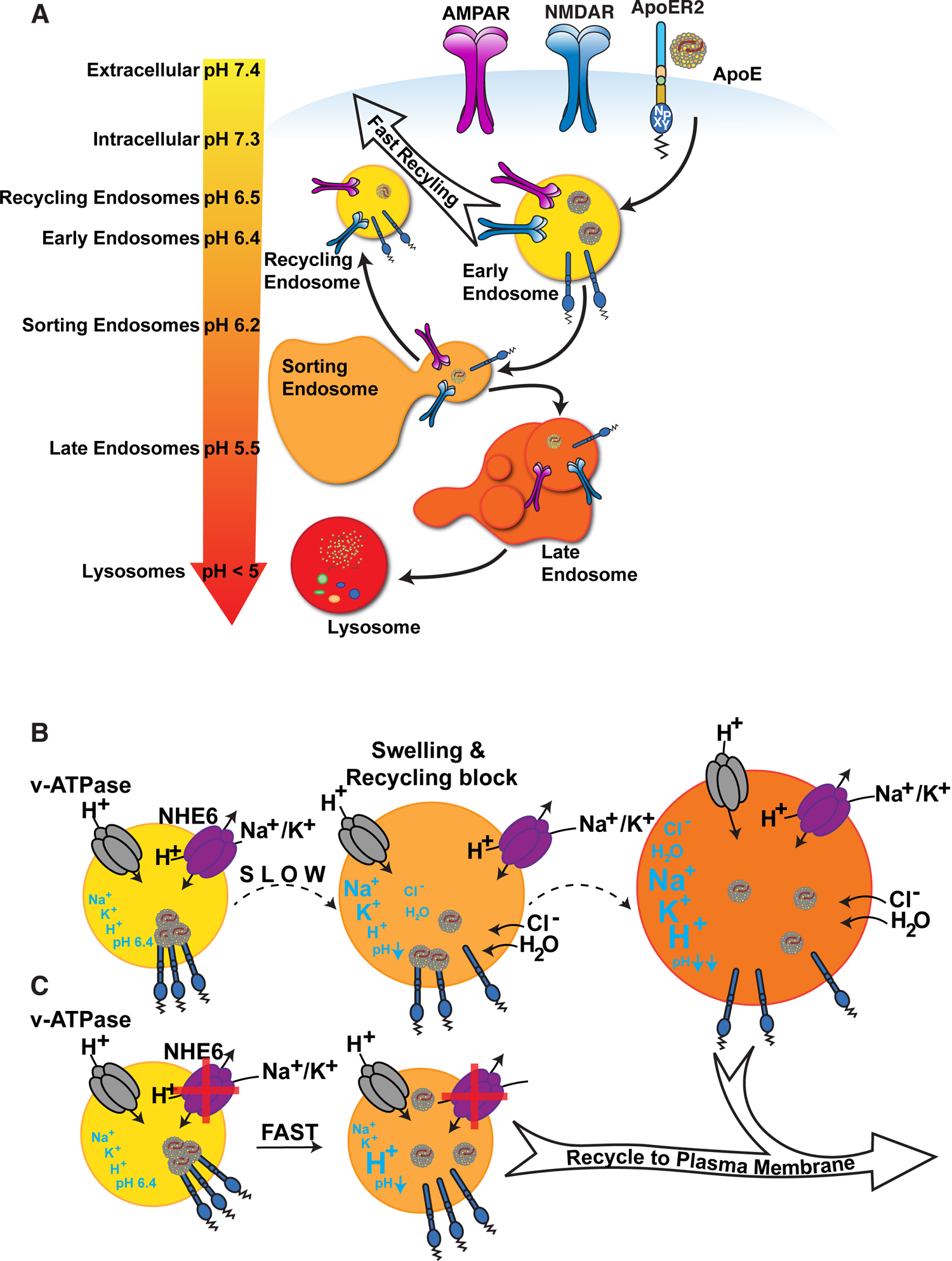
ApoE4 Induces Endolysosomal Trafficking Delay. **(A)** pH regulation within the endolysosomal pathway. Upon receptor binding ApoE is endocytosed along with glutamate receptors (AMPA and NMDA receptors). Cargo that has entered the early endosome (EE) can undergo recycling through a fast direct route without further acidification (fast recycling) or through slower sorting pathways that require further acidification (Casey et al., 2010; Naslavsky and Caplan, 2018). While lipid components are targeted to the lysosome, he majority of receptors, as well as ApoE, remain in endosomal compartments at the cellular periphery where they rapidly move back to the surface (Heeren et al., 1999). The increasingly acidic luminal pH is illustrated as a color gradient and depicted on the left. **(B)** In the presence of ApoE4 early endosomal trafficking and fast recycling are delayed. At the pH of the EE, ApoE4 is near its isoelectric point where solubility is reduced (Wintersteiner and Abramson, 1933), impairing receptor dissociation and resulting in delayed endosomal maturation with a concomitant entrapment of co-endocytosed glutamate receptors. Endosomal pH is regulated by the vesicular ATPase and the counterregulatory action of the proton leakage channel NHE6. NHE6 is an antiporter that exchanges a Na+ or K+ ion for each proton. As the pH decreases, ligands dissociate from their receptors allowing the EE to mature. If dissociation is delayed, as in case of ApoE4, endosomal trafficking is arrested, leading to progressive acidification as Na+, K+ and Cl-ions continue to enter the endosome to maintain charge neutrality while also drawing in water molecules due to osmotic pressure. We thus propose a model in which delayed ApoE4-receptor dissociation prevents early endosomal maturation and causes osmotic swelling while the pH continues to decrease until dissociation occurs. **(C)** Accelerated endosomal acidification by inhibition of the proton leak channel NHE6 resolves ApoE4 accumulation, promotes rapid receptor dissociation and promotes the vesicle entry into the lysosomal delivery or recycling pathways.

The pH of EE compartments is controlled by the vATPase-dependent proton pump and proton leakage through NHE6 (Nakamura et al., 2005). We showed that EE acidification by pharmaceutical pan-NHE inhibition or selective NHE6 knockdown in neurons prevents the ApoE4-caused trafficking delay of ApoE and glutamate receptors (Xian et al., 2018). NHE6-deficiency in humans causes neurodevelopmental defects, which result in Christianson syndrome, an X-linked genetic disorder characterized by cognitive dysfunction, autism, ataxia, and epilepsy. However, some NHE6 mutant variants causing Christianson syndrome in humans do not significantly alter the ion exchange properties of NHE6 (Ilie et al., 2020; Ilie et al., 2019) suggesting that Christianson syndrome could be caused by loss of NHE6 scaffolding functions and not by loss of endosomal pH regulation. To investigate whether NHE6-depletion can reverse ApoE4 pathology *in vivo*, we generated a conditional NHE6 knockout mouse line (NHE6-cKO) to avoid complications caused by neurodevelopmental defects by temporally and spatially controlling NHE6-ablation. We show that genetic NHE6-ablation attenuates both, the ApoE4 induced Reelin resistance and impaired synaptic plasticity in ApoE4 targeted replacement (ApoE4-KI) mice using hippocampal field recordings.

The pathological hallmarks of AD are extracellular aggregates of the amyloid β (Aβ) peptide and intracellular tangles of hyperphosphorylated tau protein. Processing of the transmembrane amyloid precursor protein (APP) at the β- and γ-sites leads to Aβ production. Aβ forms neurotoxic oligomers and accumulates in plaques. The β-site amyloid precursor protein cleaving enzyme 1 (BACE1) cleaves APP in its extracellular juxtamembrane domain to create a membrane-anchored C-terminal fragment (β-CTF) and a soluble extracellular APP domain (sAPPβ). βCTF is further cleaved by the γ-secretase complex, which leads to the release of the Aβ-peptide. APP and its secretases co-localize in endosomal compartments where cleavage can occur (Wang et al., 2018). It has further been reported that BACE1 activity is preferentially active in acidic environments (Shimizu et al., 2008). We therefore investigated whether NHE6-depletion alters BACE1 activity in neurons and whether NHE6-deficiency leads to changes in plaque deposition *in vivo*. We found that NHE6 inhibition or knockdown did not alter BACE1 activity, as judged by unchanged Aβ generation. By contrast, NHE6 ablation led to glial activation and decreased plaque load in ApoE4-KI (Sullivan et al., 1997) and APP^NL-F^ (Saito et al., 2014) double knockin mice.

### EXPERIMENTAL PROCEDURES

#### Animals

All animal procedures were performed according to the approved guidelines for Institutional Animal Care and Use Committee (IACUC) at the University of Texas Southwestern Medical Center at Dallas.

The mouse lines Rosa-stop-tdTomato B6.Cg-Gt(ROSA)26Sor^tm9(CAG-tdTomato)Hze^/J (Madisen et al., 2010) (JAX #007909) and CAG-Cre^ERT2^ B6.Cg-Tg(CAG-cre/Esr1)5Amc/J mice (Hayashi and McMahon, 2002)(JAX #004682), were obtained from The Jackson Laboratories (Bar Harbor, ME). ApoE3 and ApoE4 targeted replacement mice (ApoE3-KI, ApoE4-KI) (Knouff et al., 1999; Sullivan et al., 1997) were kind gifts of Dr. Nobuyo Maeda. APPswe (Tg2576) were generated by (Hsiao et al., 1996). The Meox-Cre B6.129S4-Meox2tm1(cre)Sor/J mice (JAX 003755) were provided by Drs. M. Tallquist and P. Soriano (Tallquist and Soriano, 2000). Conditional NHE6 knockout (NHE6-cKO) and germline NHE6 knockout (NHE6-KO) mice were generated as described below. The human APP knockin line (APP^NL-F^) (Saito et al., 2014) has been described earlier. Pregnant female SD (Sprague Dawley) rats were obtained from Charles River (SC:400). Mice were group-housed in a standard 12-h light/dark cycle and fed ad libitum standard mouse chow (Envigo, Teklad 2016 diet as standard and Teklad 2018 diet for breeding cages).

To generate NHE6-floxed (NHE6^fl^) mice, the first exon of NHE6 was flanked with loxP sites (Supplemental Fig. S1A). A loxP site was inserted 2 kb upstream of the first exon of the X-chromosomal NHE6 gene and a Neo-cassette (flanked with loxP and FRT sites) was inserted 1 kb downstream of the first exon. Insertion sites were chosen based on low conservation (mVISTA) between mammalian species (rat, human, mouse). To create the targeting vector for the NHE6-floxed mouse line, pJB1 (Braybrooke et al., 2000) was used as backbone. Murine C57Bl/6J embryonic stem (ES) cell DNA was used as template to amplify the short arm of homology (SA; kb of the first intron starting 1kb 3’ downstream of exon 1), which was inserted between the Neo and HSVTK selection marker genes of pJB1 (BamHI and XhoI sites) to create an intermediate plasmid referred to as pJB1-NHE6SA. To create the intermediate plasmid pNHE6-LA for the long homology arm, a fragment spanning from 13kb 5’ upstream to 1kb 3’ downstream of the first NHE6 exon was integrated into pCR4-TOPO (Thermo Fisher) by using a bacterial artificial chromosome (BAC, RP23 364F14, Children’s Hospital Oakland Research Institute (CHORI)) and the GAP repair technique (Lee et al., 2001) (primers to amplify the upstream (US) and downstream (DS) homology boxes are listed in the Key Resources Table). In a parallel cloning step a 2.4 kb XhoI-EcoRV NHE6-promoter region fragment spanning from 2.7 kb to 0.4 kp 5’ upstream of the NHE6 start codon was modified with the 1^st^ loxP site to generate pLoxP: three fragments (1) 0.7 kb 5’ loxP flanking NHE6-fragment, (2) 1.7 kb 3’ loxP flanking NHE6-fragment, and (3) 100 bp loxP oligo (primers/oligos for each fragment are listed in the Key Resources Table) were cloned into pCR4-TOPO. The 2.5 kb loxP-modified XhoI-EcoRV NHE6-promoter fragment of pLoxP was cloned into pNHE6-LA to create pNHE6-LA-LoxP. To obtain the final targeting construct pJB-NHE6-TV, the NotI-EagI fragment of pNHE6-LA-LoxP containing the long arm (11 kb 5’ upstream of the 1^st^ LoxP) and the 1^st^ loxP site was cloned into the NotI-site of pJB1-NHE6SA and checked for orientation (pJB1-NHE6 targeting vector). pJB-NHE6-TV was linearized with NotI and electroporated into C57Bl/6J ES cells. Gene targeting-positive C57Bl/6J ES cells (PCR-screen) were injected into albino C57Bl/6J blastocysts, resulting in chimeric mice. The chimeras were crossed to C57Bl/6J mice, resulting in NHE6^fl/+^ females. Genotyping: The NHE6-floxed PCR amplifies a 250 bp of the wildtype and 270 bp of the floxed allele, primers are listed in the Key Resources Table.

To generate NHE6-cKO, NHE6^fl/+^ females were crossed to CAG-Cre^ERT2^ mice to obtain NHE6^fl/fl^;CAG-Cre^ERT2^ and NHE6^y/fl^;CAG-Cre^ERT2^ mice (NHE6-cKO) and CAG-Cre^ERT2^ negative control littermates (NHE6-floxed). NHE6-cKO mice were backcrossed to ApoE3-KI or ApoE4-KI mice. Breeding pairs were set up in which only one parent was CAG-Cre^ERT2^ positive. The following genotypes were used for hippocampal field recordings: (1) ApoE3^ki/ki^;NHE6^y/fl^ (short: ApoE3-KI), (2) ApoE3^ki/ki^;NHE6^y/fl^;Cre^ERT2^ (short: ApoE3-KI;NHE6-cKO), (3) ApoE4^ki/ki^;NHE6^y/fl^ (short: ApoE4-KI), and (4) ApoE4^ki/ki^;NHE6^y/fl^;Cre^ERT2^ (short: ApoE4-KI;NHE6-cKO). The ApoE4-KI;NHE6-cKO line was further crossed with the APP^NL-F^ line to generate ApoE4-KI;NHE6-cKO;APP^NL-F^ and ApoE4-KI;NHE6-floxed;APP^NL-F^ control littermates. To induce genetic depletion of NHE6, tamoxifen (120 mg/kg) was intraperitoneally injected at 6-8 weeks of age. Injections were applied for five consecutive days. Tamoxifen was dissolved in sunflower oil (Sigma, W530285) and 10% EtOH.

To generate the germline NHE6 knockout line (NHE6-KO), heterozygous NHE6^fl/+^ females were crossed to Meox-Cre to yield germline mutant NHE6^-/-^ females and NHE6^y/-^ males (NHE6-KO). Genotyping: The NHE6-floxed PCR amplifies 250 bp of the wildtype, 270 bp of the floxed, and no fragment in the knockout alleles. Recombination was verified with the NHE6-rec PCR, which amplifies 400 bp if recombination has occurred. PCR-primers are listed in the Key Resources Table. NHE6-KO animals were further crossed with APP^NL-F^ (Saito et al., 2014) mice. APP^NL-F^;NHE6-KO (APP^NL-F/NL-F^;NHE6^y/-^) and control (APP^NL-F^ = APP^NL-F/NL-F^;NHE6^y/+^) littermates were obtained by crossing NHE6^+/-^;APP^NL-F/NL-F^ females with NHE6^y/-^;APP^NL-F/NL-F^ males.

### DNA Constructs, Recombinant Proteins, Lentivirus Production

Lentiviral plasmids with shRNA against NHE6 and the scrambled control have been described in Xian et al.(Xian et al., 2018). Plasmids encoding ApoE3 and ApoE4 (Chen et al., 2010), and Reelin (D’Arcangelo et al., 1997) have been described before. The lentiviral plasmid encoding the Vamp3-pHluorin2 fusion protein was cloned by inserting mouse Vamp3 cDNA, a linker and pHluorin2 (pME, Addgene #73794) (Stawicki et al., 2014) into pLVCMVfull (Xian et al., 2018). For Vamp3 the for primer (5’-TTCAAGCTTCACCATGTCTACAGGTGTGCCTTCGGGGTC-3’) contains a Kozak sequence, the reverse primer encodes a KLSNSAVDGTAGPGSIAT-linker (Nakamura et al., 2005) (5’ CATTGTCATCATCATCATCGTGTGGTGTGTCTCTAAGCTGAGCAACAGCGCCGTGGACGGCACCGCCGGCCCCGGCAGCATCGCCACCAAGCTTAAC-3’). The pHluorin2 primers were for 5’-CCGGTCCCAAGCTTATGGTGAGCAAGGGCGAGGAGCTGTTC-3’ and rev 5’-GCCCTCTTCTAGAGAATTCACTTGTACAGCTCGTCCATGCCGTG-3’. The fragments were sequentially cloned into pcDNA3.1 and the fusion protein was then transferred with NheI and EcoRI into pLVCMVfull. Lentiviral plasmids psPAX2 and pMD2.g were a kind gift from Dr D. Trono and obtained from Addgene.

Recombinant Reelin and ApoE were generated in 293HEK cells. Reelin was purified as described before (Weeber et al., 2002). ApoE-conditioned medium was collected from HEK293 cell cultures transiently transfected with pcDNA3.1-ApoE constructs or empty control vector (pcDNA3.1-Zeo). ApoE concentration was measured as described before (Xian et al., 2018).

For lentivirus production HEK 293-T cells were co-transfected with psPAX2, pMD2.g, and the individual shRNA encoding transfer or Vamp3-pHluorin2 constructs. Media was replaced after 12-16 hours. Viral particle containing media was collected after centrifuging cellular debris. The viral particles were 10x concentrated by ultra-centrifugation and resuspended in Neurobasal medium. To infect neurons on DIV7 1/10^th^ of the culture medium was replaced by concentrated virus. After 24 hours the virus was removed.

### Histochemistry

Mice were euthanized with isoflurane and perfused with PBS followed by 4% paraformaldehyde (PFA) in PBS. Brains were removed and post-fixed for 24 hours in 4% PFA. Fresh fixed brains were immobilized in 5% agarose in PBS and 50 μm thick sections were sliced on a vibratome (Leica, Wetzlar, Germany). Vibratome slices of Rosa-stop-tdTomato;CAGCre^ERT2^ mice, with or without tamoxifen injection at 8 weeks of age were mounted with Antifade Mounting medium containing DAPI (Vectashield). For immunofluorescence, vibratome slices were labeled for Calbindin after permeabilization with 0.3% Triton X in PBS and blocking for 1 hour in blocking buffer (10% normal goat serum, 3% BSA, and 0.3% Triton X in PBS). The primary antibody mouse anti-Calbindin (Swant CB300) was diluted in blocking buffer (1:1000) and added to the slices for 24-48 h at 4°C. Slices were subsequently washed 4x 15 min with PBS containing 0.3% Triton X. Slices were incubated with anti-mouse IgG coupled to Alexa594 (1:500 in blocking buffer) for 2 hours at room temperature. After washing, slices were mounted with Antifade Mounting medium with DAPI (Vectashield). Images were taken with an Axioplan 2 microscope (Zeiss).

For immunohistochemistry staining, PFA fixed brains were block-sectioned into coronal slabs, paraffin-embedded, and serially sectioned on a rotating microtome (Leica) at 5 μm. Deparaffinized sections were stained with Thioflavin S as described before (Guntern et al., 1992). Briefly, deparaffinized slices were oxidized in 0.25% KMnO_4_ for 20 min. After washing with water slices were bleached with 1% K_2_S_2_O_5_ / C_2_H_2_O_4_ for 2 min, washed in water, and treated with 2% NaOH/H_2_O_2_ for 20 min. After washing with water slices were acidified in 0.25% CH_3_COOH for 1 min, washed with water and equilibrated in 50% EtOH for 2 min, and stained in Thioflavin S solution for 7 min. Reaction was stopped by washing in 50% EtOH. Slices were dehydrated with 95% EtOH, followed by 100% EtOH and Xylene. Slices were mounted with Cytoseal 60 (Thermo Scientific). Deparaffinized sections were labeled using antibodies raised against GFAP (Abcam), Iba1 (Waco), and Aβ (4G8, Covance or 6E10, Biolegend). Briefly, 5 μm sections were deparaffinized, subjected to microwave antigen retrieval (citrate buffer, pH 6.0), permeabilized with 0.3% (vol/vol) Triton X, endogenous peroxidases activity was quenched for Diaminobenzidine (DAB) staining. Slices were blocked with goat serum (2.5 %) prior to overnight incubation with primary antibodies at 4°C. Primary antibodies were detected by either fluorescent secondary antibodies (goat-anti-mouse Alexa594, goat-anti-rabbit Alexa488) or sequential incubation with biotinylated secondary antisera and streptavidin-peroxidase for DAB staining. Diaminobenzidine chromagen was used to detect the immunoperoxidase signal (Sinclair et al., 1981) (Vector; anti-mouse and anti-rabbit IgG kits). Fluorescence-labeled slices were counter-stained with DAPI (mounting media with DAPI, Vectashield). Standard protocols were utilized for staining of paraffin sections with hematoxylin and eosin (H&E; Leica) (Fischer et al., 2008). Microscopy was performed on a high-throughput microscope (NanoZoomer 2.0-HT, Hamamatsu) for DAB stained tissue or with an Axioplan 2 microscope (Zeiss) for immunofluorescence analysis. The analysis of DAB labeled antibodies was performed after exporting the images with NDP.view2 software with ImageJ. For Thioflavin S staining and Aβ labeling plaques were quantified by categorizing them as small, medium, and large (Thioflavin S) or dense and diffuse (4G8) as depicted in Supplemental Figure S4. Co-localization analysis of microglia (Iba1) and astrocytes (GFAP) with plaques (6E10) was performed with ImageJ. A blind observer selected the area of plaques with circles of 20, 40, or 80 µm diameter and analyzed the intensity of 6E10 and Iba1 or GFAP by using the ImageJ plugin RGB_measure. 6E10 positive microglia (Iba1) were quantified by a blind observer by first identifying microglia structures/cells in the green channel (Iba1), and then analyzing the proportion of 6E10 positive structures and signal intensity (red channel).

### Primary Cortical Neuronal Cultures

Primary cortical neuronal cultures were prepared from rat (SD, Charles River) or various mouse lines (wildtype, NHE6-KO, APPswe) (E18) as described previously (Chen et al., 2005). Neurons were cultured in Poly-D-Lysine coated 6-well plates (1 million / 9 cm^2^) or on coverslips (30,000 neurons / 1.1 cm^2^) in presence of Neurobasal medium supplemented with B27, glutamine, and Penicillin-Streptomycin at 37°C and 5% CO_2_. At indicated days in vitro (DIV) primary neurons were used for experiments.

### Mouse Embryonic Fibroblasts

Fibroblasts were isolated from NHE6-KO and wildtype littermate control embryos (E13.5). After removing the head and the liver, the tissue was trypsinized (0.05% trypsin-EDTA) at 4°C overnight, followed by 30 min at 37°C. Suspended cells were cultivated in DMEM high glucose with 15% FCS, 2 mM L-glutamine, and Pen/Strep. Fibroblasts were serially passaged until proliferation slowed down. Immortalization was achieved by keeping fibroblasts in culture under high confluency until they overcame their growth-crisis.

### pH Measurements

Mouse embryonic fibroblasts derived from either NHE6-KO or wildtype littermate embryos were infected with Vamp3-pHluorin2 lentivirus. 24-48 hours post infection, vesicular pH was measured on a Zeiss LSM880 Airyscan confocal microscope as described in Ma et al. (Ma et al., 2017). For the standard calibration curve, cells were washed and incubated with pH standard curve buffer (125 mM KCl, 25 mM NaCl, 10 μM monensin, 25 mM HEPES for pH > 7.0 or 25 mM MES for pH < 7.0; pH adjusted with NaOH and HCl) and imaged in 5% CO_2_ at 37°C. For vesicular pH measurements, cells were washed and imaged with pH standard curve at pH 7 without monensin. Samples were excited at 405 and 488 nm with an emission of 510 nm. The same settings were used for every image, and images were analyzed using ImageJ Software. The intensity of excitations with 405 and 488 nm was measured, individual vesicles were marked as regions of interest, and the 405/488 ratio was calculated and plotted against pH for the standard curve.

### Biochemistry

To analyze receptor recycling cell surface biotinylation was performed. At DIV10-14, primary neurons were pre-treated for 30 min with ApoE-conditioned medium (5 μg/ml) and incubated with Reelin (2 μg/ml) for an additional 30 min (see timeline in **Figure 3A**). After treatment, cells were washed with cold PBS and incubated in PBS containing sulfo-NHS-SS-biotin for 30 min at 4°C. Subsequently excess reagent was quenched by rinsing the neurons with cold PBS containing 100 mM glycine. Neurons were lysed in 160 µl/ 9 cm^2^ lysis buffer (PBS with 0.1% SDS, 1% Triton X-100, and protease inhibitors) at 4°C for 20 min. Cell debris were pelleted at 14,000 rpm for 10 min at 4°C. The protein concentration was measured using the Bradford Protein Assay (Bio-Rad). One hundred µg of total proteins were incubated with 50 µl of NeutrAvidin agarose at 4°C for 1 hour. Agarose beads were washed three times using washing buffer (500 mM NaCl; 15 mM Tris-HCl, pH 8.0; 0.5% Triton X-100), biotinylated surface proteins were eluted from agarose beads by boiling in 2x SDS sample loading buffer and loaded on an SDS-PAGE gel for Western blot analysis. GST-control and GST-RAP (50 μg/ml) pre-treatment of neurons was performed for 1 hour.

To analyze BACE1 activity, βCTF was detected by immune blotting. BACE1 activity was examined after pharmacological NHE inhibition or genetic NHE6 knockdown in primary neurons of APPswe mice. For NHE inhibition DIV10 neurons were treated with 5 µg/ml ApoE4, 3 µM EMD87580 (Merck), and/or 1 µM γ-secretase inhibitor L-685458 (Merck) for 5 hours. For knockdown of NHE6 DIV7 neurons of APPswe mice were infected with lentivirus encoding shRNA against NHE6 or a scrambled control shRNA. On DIV13 neurons were treated with γ-secretase inhibitor for 12 hours. Proteins were extracted for Western blot analysis: Cells were washed three times with cold PBS, and lysed for 20 min on ice in RIPA buffer (50 mM Tris-HCl, pH 8.0; 150 mM NaCl; 1% Nonidet P-40; phosphatase and proteinase inhibitors). Cell debris were pelleted at 14,000 rpm for 10 min at 4°C. Protein concentration was measured using the Bradford Protein Assay (Bio-Rad). After adding 4xSDS loading buffer (0.1 M Tris-HCl, pH 6.8, 2% SDS, 5% β-mercaptoethanol, 10% glycerol, and 0.05% bromophenol blue) samples were denatured at 80°C for 10 min. Proteins were separated via SDS-PAGE and transferred to a nitrocellulose membrane for Western blotting using different antibodies listed in the Key Resources Table.

Brain tissue was dissected and prepared for immunoblotting as followed: After removal, brains were placed in ice-cold PBS containing proteinase inhibitors. Anatomical sectioning was performed under a microscope. The hippocampus was dissected out and transversal slices were further separated into Cornu Ammonis (CA) 1, CA3, and dentate gyrus. Respective pieces of the same anatomical regions of one brain were pooled, shock-frozen in liquid nitrogen and stored at −80C. Frozen tissue was homogenized in 10x volume of RIPA buffer and incubated on ice for 30 min. Cell debris were pelleted for 10 min with 14,000 rpm at 4°C. After adding 4xSDS loading buffer the samples were denatured at 80°C for 10 min and used for immunoblotting.

To measure soluble and insoluble Aβ species, a sequential homogenization procedure was employed. After removal of the brains from PBS perfused mice, cortical tissue was dissected and flash frozen. Frozen cortical tissue was homogenized in TBS supplemented with phosphatase and proteinase inhibitors at 100 mg protein/ml using a glass dounce homogenizer. Crude lysate was centrifuged at 800 *xg* for 5 min at 4°C. The supernatant was further centrifuged at 100,000 *xg* for 30 min at 4°C and collected as TBS-soluble lysate (Aβ-soluble). The TBS-pellet was further homogenized in 1% Triton-TBS containing phosphatase and proteinase inhibitors, centrifuged at 100,000 *xg* for 30 min at 4°C and collected as 1% Triton-soluble lysate. The Triton-pellet was incubated with 5M guanidine-HCl rotating at RT for 1 hour. Guanidine soluble lysate (Aβ-insoluble) was collected after centrifugation at 21,000 *xg* for 15 min at 4°C. The guanidine-pellet was further solubilized in 1/20^th^ volume with 70% Formic Acid (Aβ-insoluble) and centrifuged at 21,000 *xg* for 15 min at 4°C. Soluble and insoluble Aβ levels were measured in duplicates using a commercial Aβ42 ELISA kit (ThermoFisher, KHB3441) following the manufacturer’s instructions.

### Extracellular Field Recordings

Hippocampal slices were prepared from 3-month-old mice (tamoxifen-injected at 8 weeks). Slices of mice were obtained from four different genotypes; NHE6 cKO mice or NHE6 floxed mice with ApoE3-KI or ApoE4-KI (ApoE3^ki/ki^;NHE6^y/fl^, ApoE3^ki/ki^;NHE6^y/fl^;Cre^ERT2^, ApoE4^ki/ki^;NHE6^y/fl^, or ApoE4^ki/ki^;NHE6^y/fl^;Cre^ERT2^). The brains were quickly removed and placed in ice cold high sucrose cutting solution (in mM: 110 sucrose, 60 NaCl, 3 KCl, 1.25 NaH_2_PO_4_, 28 NaHCO_3_, 0.5 CaCl_2_, 5 glucose, 0.6 Ascorbic acid, 7 MgSO_4_), bubbled with a gas mixture of 95% O_2_ and 5% CO_2_ for oxygenation. 350 µm transverse sections were cut using a vibratome (Leica). Slices were transferred into an incubation chamber containing 50% artificial cerebrospinal fluid (aCSF, in mM: 124 NaCl, 3 KCl, 1.25 NaH_2_PO_4_, 26 NaHCO_3_, 10 D-glucose, 2 CaCl_2_, 1 MgSO_4_) and 50% sucrose cutting solution, oxygenated with 95%O_2_/5%CO_2_. Slices were transferred into an oxygenated interface chamber and perfused with aCSF with or without Reelin (2 μg/ml). The stimulating electrode was placed on the Schaffer-collateral of the CA1-pyramidal neurons and the recording electrode on the dendrites of the CA3-pyramidal neurons. Once baseline was stably recorded for 20 min, theta burst was applied and traces collected for an hour. For stimulation concentric bipolar electrodes (FHC, catalog no CBBRC75) were placed into the stratum radiatum. Stimulus intensity was set at 40-60% of maximum response and delivered at 33 mHz through an Isolated Pulse Stimulator (A-M Systems, Model 2100). A custom written program in LabView 7.0 was used for recording and analysis of LTP experiments. A theta burst (TBS; train of 4 pulses at 100 Hz repeated 10 times with 200 ms intervals; repeated 5 times at 10 s intervals) was used as a conditioning stimulus.

### Statistical Methods

Data were expressed as the mean ± SEM and evaluated using two-tailed Student’s t-test for two groups with one variable tested and equal variances, one-way analysis of variance (ANOVA) with Dunnett’s post-hoc or Tukey’s post-hoc for multiple groups with only variable tested, or two-way ANOVA with Sidak’s post-hoc for plaque quantification (two independent variables of NHE6 genotype and plaque classification). The differences were considered to be significant at p<0.05 (*p<0.05, ** p<0.01, *** p<0.001). Software used for data analysis was ImageJ (NIH), LabView7.0 (National Instruments), Odyssey Imaging Systems (Li-Cor), Prism7.0 (GraphPad Software).

## RESULTS

### NHE6 is Required for Postnatal Purkinje Cell Survival

NHE6 germline knockout mice (NHE6-KO) and tamoxifen-inducible conditional NHE6 knockout mice (NHE6-cKO) were generated as described in Experimental Procedures and **Supplemental Figure S1A-C**. To validate early endosomal pH acidification by NHE6-deficiency we isolated mouse embryonic fibroblasts from the NHE6-KO line and infected them with a Vamp3-pHluorin2 lentivirus expressing a fusion protein consisting of the endosomal Vamp3 protein and the ratiometric pH indicator pHluorin2 (Stawicki et al., 2014). We found a significantly reduced number of vesicles with pH 6.4 and above in NHE6-KO fibroblasts when compared to control (**Supplemental Figure S1D-F**).

To induce the conditional KO of NHE6, tamoxifen was administered to NHE6-cKO (NHE6-floxed;CAG-Cre^ERT2^) mice at two months (**Figure 2A**) and experiments were performed at the indicated time points. NHE6 knockout efficiency in the brains of tamoxifen injected NHE6-cKO mice was 65-82% and varied between brain regions (**Figure 2B**). To further investigate Cre^ERT2^ activity upon tamoxifen-injection, we bred the CAG-Cre^ERT2^-line with a stop-tdTomato reporter line in which a floxed stop-codon precedes the tdTomato start-codon. After tamoxifen injection, brains were examined for tdTomato expression. Without tamoxifen injection, tdTomato expressing cells were almost absent in the hippocampus. Tamoxifen induced recombination led to a broad expression of tdTomato in the hippocampus **(Supplemental Figure S1G**).

**Figure 2.**
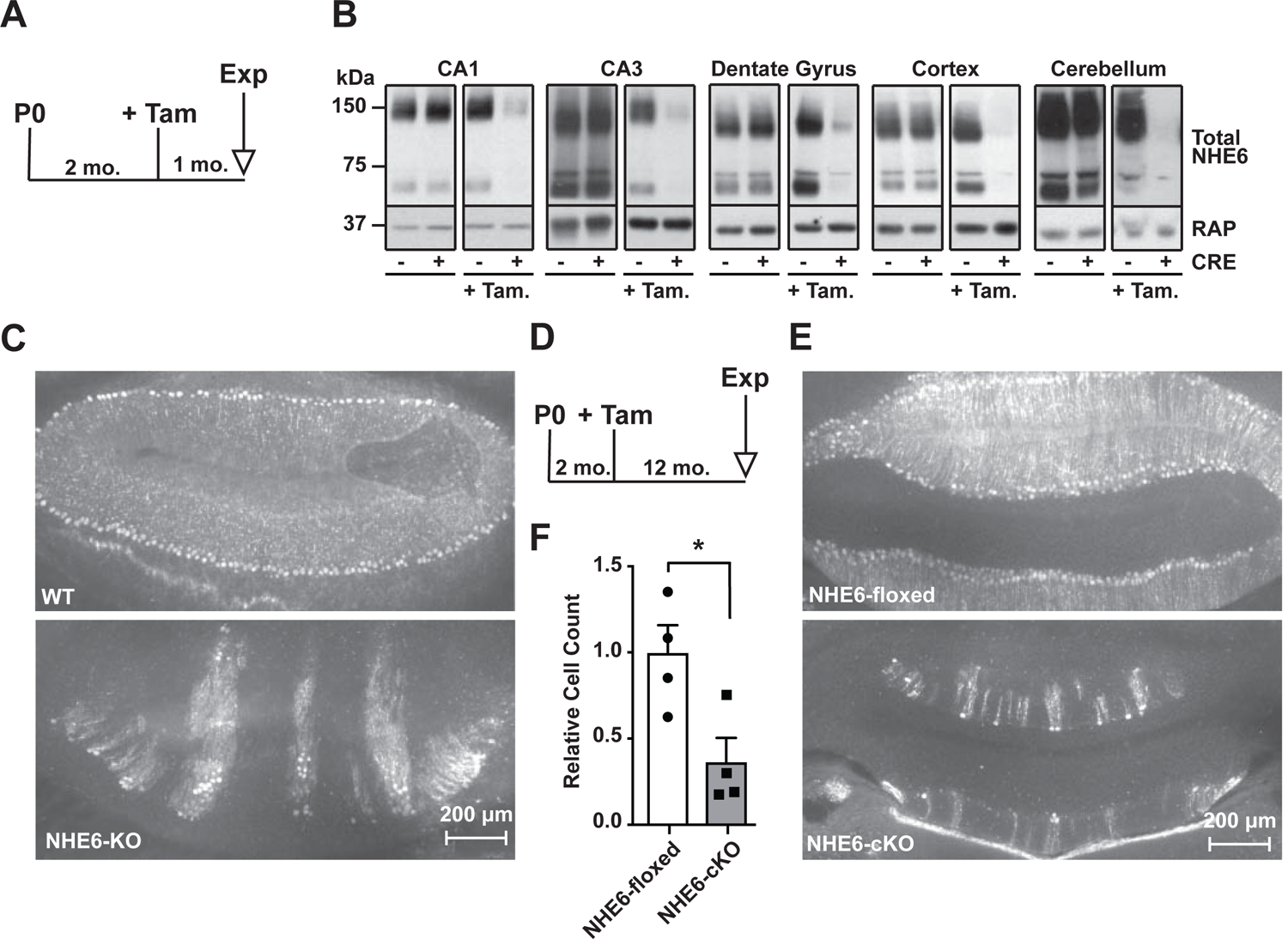
Long-Term NHE6-Deficiency Induced After Purkinje Cell Maturation Causes Purkinje Cell-loss. **(A)** Experimental timeline for B, mice were injected with tamoxifen at 2 months; after one month the brains were analyzed for NHE6 expression (Tam = tamoxifen, Exp. = experiment, mo. = months). **(B)** Western blot showing the efficiency of tamoxifen-induced NHE6 knockout in different brain regions (CA1, CA3, dentate gyrus, cortex, and cerebellum). The knockout efficiency differed between brain regions, it was 80±2% in CA1, 82±5.7% in the CA3, 67±6.8% in the dentate gyrus, 65±11.2% in the cortex, and 74±4.7% in the cerebellum. **(C-F)** NHE6 deficiency leads to cerebellar Purkinje cell loss in germline (NHE6-KO, C) and conditional (NHE6-cKO, D-F) knockout mice. The timeline shows that NHE6-cKO and control mice were tamoxifen-injected at two months and analyzed one year after (D). Calbindin was fluorescently labeled to highlight Purkinje cells in the cerebellum. Massive loss of Purkinje cells was found in NHE6-KO (C, lower panel), compared to wildtype control (C, upper panel). Long-term loss of NHE6, induced after Purkinje cell maturation at two months of age, also led to massive Purkinje cells-loss when mice were examined one year after NHE6-ablation (E, lower panel). **(F)** Quantification of Purkinje cell-loss in the cerebellum of NHE6-cKO mice. NHE6-flox mice represent Cre^ERT2^-deficient, tamoxifen injected mice. Values are expressed as mean ± SEM from 4 independent experiments. Statistical analysis was performed using Student *t*-test. *p<0.05.

Individuals with Christianson Syndrome and mice lacking NHE6 present with motor deficits due to a dramatic progressive loss of cerebellar Purkinje cells (Ouyang et al., 2013). We have reproduced the Purkinje cell loss in our germline NHE6-KO mice (**Figure 2C**). Next, we investigated whether Purkinje cell loss is the consequence of neurodevelopmental or neurodegenerative effects caused by loss of NHE6. NHE6-deficiency was induced at 2 months, after Purkinje cells had developed and matured. One year after NHE6-ablation Purkinje cell loss was indistinguishable from that seen in the germline knockouts (**Figure 2D-F**). Therefore, Purkinje cell degeneration manifests itself postnatally and is not developmentally determined by the absence of NHE6. However, it is possible that loss of scaffolding functions and proper sorting, rather than dysregulation of endosomal pH, could be the main mechanism that causes Christianson syndrome, including Purkinje cell loss (Ilie et al., 2020; Ilie et al., 2019). If this could be substantiated by the development or discovery of NHE6 mutants that selectively ablate its ion exchange capacity without affecting its subcellular sorting or interaction with cytoplasmic or luminal binding partners, this would further raise the potential of NHE6 as a novel drug target for neurodegenerative diseases.

### Genetic Disruption of NHE6 Restores Trafficking of Apoer2, AMPA and NMDA Receptors in the Presence of ApoE4

As we reported previously, ApoE4 impairs the trafficking of synaptic surface receptors (Chen et al., 2010). To monitor receptor recycling in neurons we made use of an assay where Reelin is used to modulate receptor surface expression. Reelin is applied to primary neurons for 30 minutes in the presence or absence of naturally secreted, receptor binding-competent ApoE (**Figure 3**). Subsequently, surface biotinylation is performed and cells are harvested for immunoblotting to quantify the amount of Apoer2 and glutamate receptors expressed on the neuronal surface (Chen et al., 2010; Xian et al., 2018).

We have shown previously that in the presence of ApoE4, Apoer2 and glutamate receptors recycle poorly to the neuronal surface. This recycling block could be resolved by endosomal acidification induced by shRNA knockdown of NHE6 or by applying the NHE inhibitor EMD87580 (Xian et al., 2018). To further exclude a nonspecific effect caused by the inhibition of other NHE family members or by shRNA off-target effects, we applied this assay on neurons isolated from NHE6-KO embryos (**Figure 3B**). Apoer2 recycling was completely restored in NHE6-KO neurons treated with ApoE4 (**Figure 3C**). We previously reported that the addition of ApoE3 to neurons also affects Apoer2 trafficking to a small, but reproducible extent. This was also abolished in NHE6-KO neurons (**Figure 3C**). In addition, genetic loss of NHE6 equally restored the ApoE4-impaired surface expression of AMPA and NMDA receptor subunits (**Figure 3D-F**).

**Figure 3.**
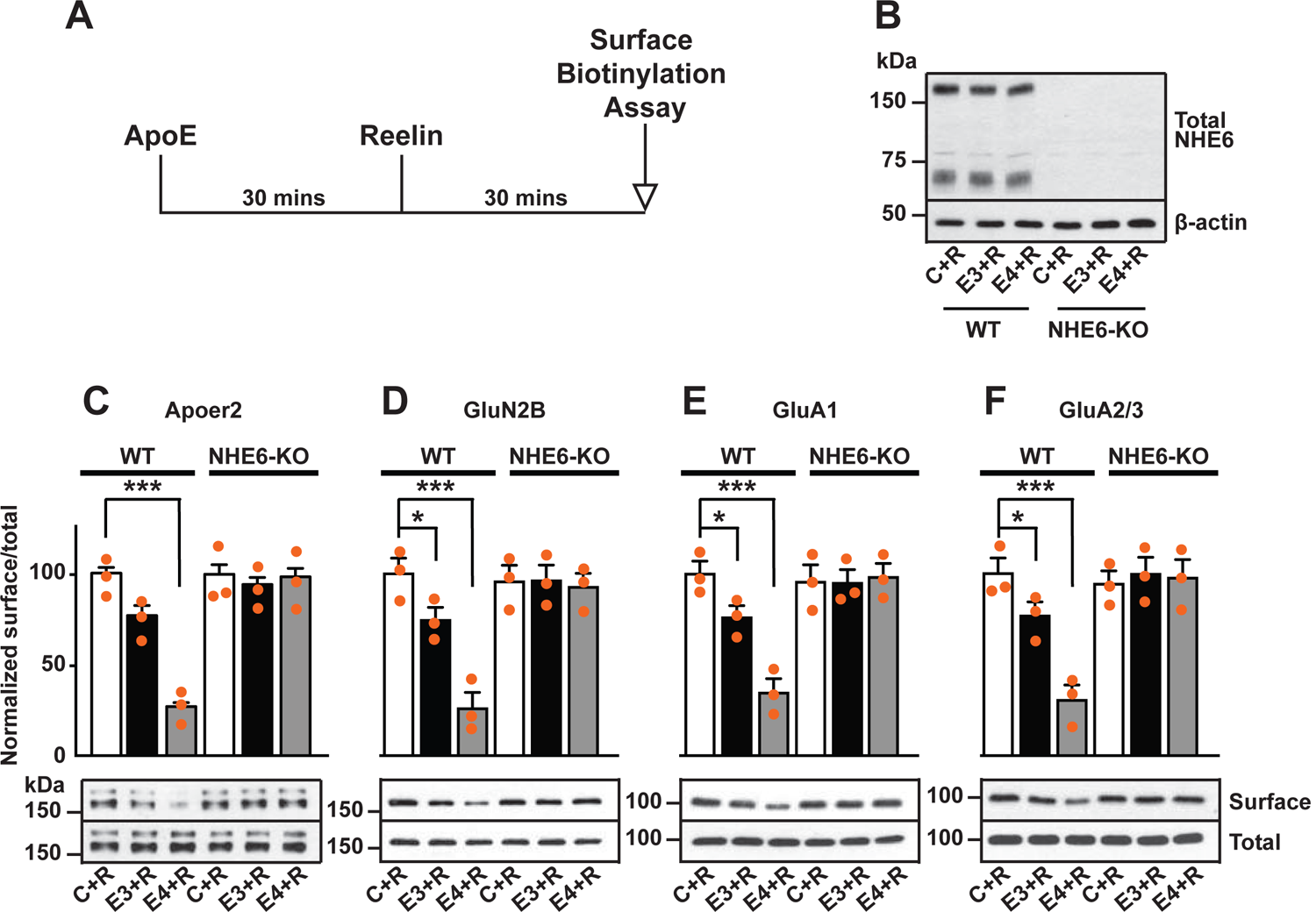
NHE6 Knockout Alleviates ApoE4-Impaired Surface Trafficking Deficits of Apoer2 and Glutamate Receptors. **(A)** Timeline for the receptor surface expression assay applied for the experiments shown in B-F. Primary neurons were treated with naturally secreted ApoE3 or ApoE4 and/or Reelin before they underwent surface biotinylation. **(B-F)** Wildtype and NHE6-KO primary neurons were prepared from littermates and used in the receptor surface expression assay described in A. **(B)** NHE6-deficiency was confirmed via Western blot, β-actin was used as loading control. **(C-F)** ApoE-conditioned media treatment reduces the surface expression of Apoer2 and glutamate receptors in presence of Reelin in primary neurons. Receptor surface levels show a stronger reduction with ApoE4 than ApoE3. NHE6 depletion counteracts the ApoE4-induced reduction of receptor surface expression. Cell surface biotinylation assay was performed for Apoer2 (C), GluN2B (D), GluA1 (E) and GluA2/3 (F). Total levels were analyzed by immunoblotting of whole cell lysates against the same antibodies. β-actin was used as loading control. Quantitative analysis of immunoblot signals is shown in the lower panels (C-F). All data are expressed as mean ± SEM from 3 independent experiments. *p < 0.05, **p < 0.01, ***p < 0.005. Statistical analysis was performed using one-way ANOVA and Dunnett’s post hoc test **(C-F)**.

### Conditional Disruption of NHE6 Relieves Synaptic Reelin Resistance in ApoE4-KI mice

Reelin can enhance long term potentiation (LTP) in hippocampal field recordings of ApoE3-KI but not ApoE4-KI acute brain slices (Chen et al., 2010). We previously showed that this Reelin-resistance in ApoE4-KI slices was attenuated by pharmacological NHE inhibition (Xian et al., 2018). To investigate if endogenous loss of NHE6 also restores synaptic plasticity in the presence of ApoE4 we performed hippocampal field recordings on NHE6-cKO mice bred to ApoE3-KI or ApoE4-KI mice. To avoid potentially compounding effects of NHE6 deficiency during embryonic development (Ouyang et al., 2013), NHE6 gene disruption was induced at 8 weeks by intraperitoneal tamoxifen injection (Lane-Donovan et al., 2015). Electrophysiology was performed 3-4 weeks after NHE6-depletion in 3 months old mice (NHE6-cKO, **Figure 4B, 4D, 4F, and 4H**). Tamoxifen-injected NHE6-floxed Cre^ERT2^-negative mice expressing human ApoE3 or ApoE4 served as controls (ApoE3-KI and ApoE4-KI, **Figure 4A, 4C, 4E, and 4G**). For field recordings hippocampal slices were perfused with Reelin as described (Beffert et al., 2005; Chen et al., 2010; Durakoglugil et al., 2009; Weeber et al., 2002). Conditional genetic loss of NHE6 resulted in a moderate reduction of the ability of Reelin to enhance LTP in ApoE3-KI mice (comparing **Figure 4A and 4E to 4B and 4F**). By contrast, as reported previously (Chen et al., 2010), hippocampal slices from ApoE4-KI mice were completely resistant to LTP enhancement by Reelin **(Figure 4C and 4G**). This resistance was abolished when NHE6 was genetically disrupted in ApoE4-KI mice: Reelin application enhanced LTP (**Figure 4D and 4H**) in ApoE4-KI;NHE6-cKO to a comparable extent as in the ApoE3-KI;NHE6-cKO mice (**Figure 4B**). Synaptic transmission was monitored and input-output curves were generated by plotting the fiber volley amplitude, measured at increasing stimulus intensities, against the fEPSP slope. No significant differences were found (**Figure 4I and 4J**).

**Figure 4.**
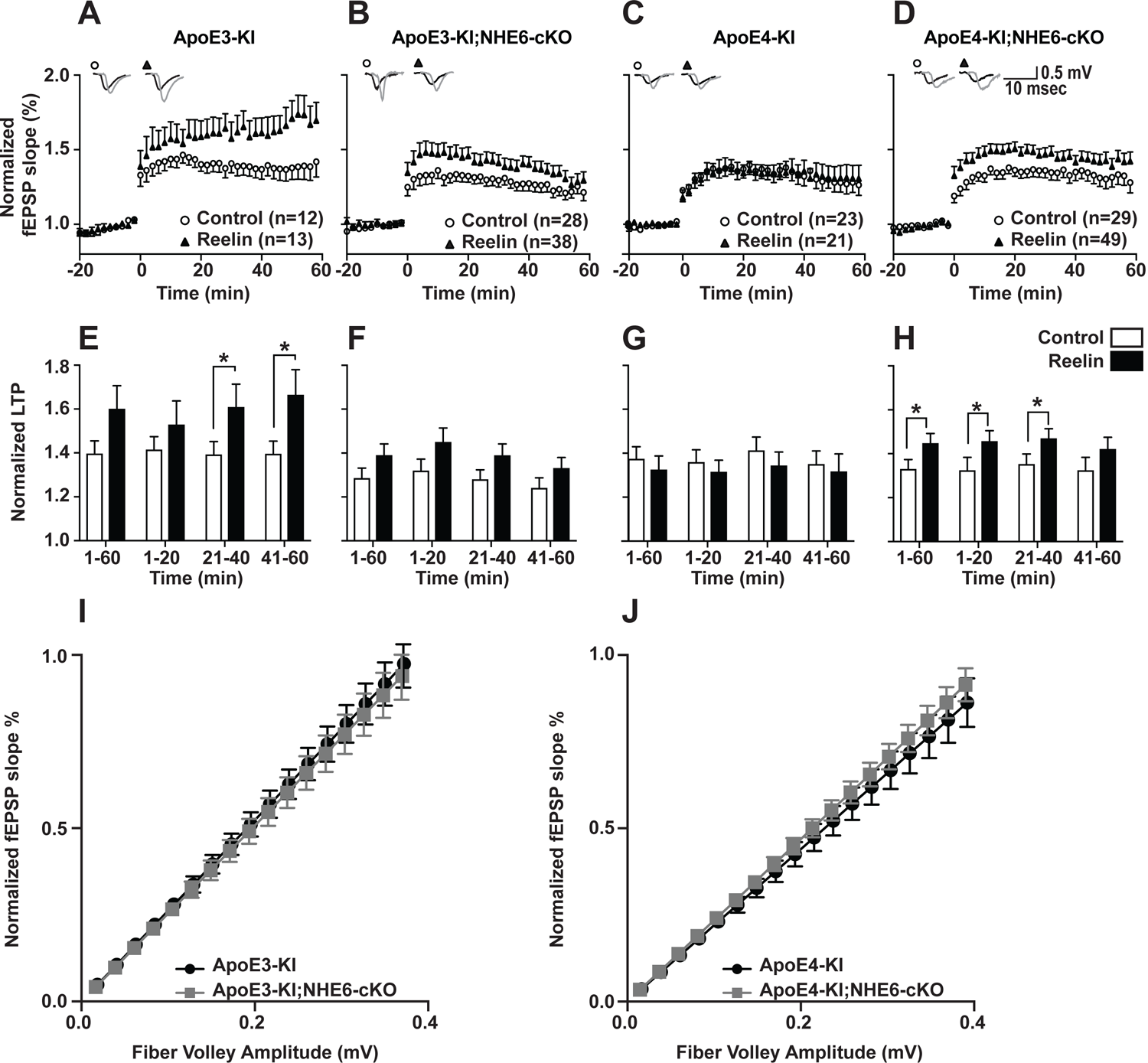
Effect of Conditional NHE6 Knockout on Reelin-potentiated Synaptic Plasticity. **(A-H)** Conditional knockout of NHE6 in ApoE4-KI mice attenuates Reelin-enhanced long-term potentiation (LTP). Reelin facilitated induction of LTP in ApoE3-KI **(A, E)**, but not ApoE4-KI **(C, G)** control (NHE6-floxed) mice. NHE6-deficiency in ApoE3-KI mice caused a reduction in Reelin enhanced LTP, such that it is not significantly different from the control LTP **(B, F)**. Importantly, in ApoE4-KI;NHE6-cKO mice Reelin was able to enhance theta-burst induced potentiation **(D, H)**. Hippocampal slices were prepared from 3 months old double mutant mice with either human ApoE3-KI or ApoE4-KI crossed with NHE6 conditional knockout mice (NHE6-cKO, tamoxifen-injections at 6-8 weeks). Extracellular field recordings were performed in slices treated with or without Reelin. Theta burst stimulation (TBS) was performed after 20 minutes of stable baseline. Representative traces are shown in each panel, before TBS induction (black) and 40 min after TBS (grey). **(E-H)** Quantitative analysis of normalized fEPSP slopes at time intervals as indicated. **(I, J)** Input output curves of ApoE3-KI **(I)** and ApoE4-KI **(J)** mice with or without NHE6-cKO. All data are expressed as mean ± SEM. N-numbers for each genotype group and treatment are indicated in panels A-D. *p < 0.05. Statistical analysis was performed using Student *t*-test.

### NHE Inhibition or NHE6 Knockdown does not Alter **β**-CTF Generation *in vitro*

Cleavage of APP by γ-secretase and the β-site APP cleaving enzyme 1 (BACE1) generates the short neurotoxic polypeptide Aβ. Cleavage by BACE1 results in a membrane anchored fragment called βCTF, which is further processed by γ-secretase to yield the soluble Aβ peptide. BACE1 processing of APP occurs in the Golgi complex, on the cell membrane, and after endocytosis in endosomes (Caporaso et al., 1994; Vassar et al., 1999). It has been reported that BACE1 activity increases with lower pH (Hook et al., 2002). In a recent *in vitro* study in a HEK293 cell line overexpressing APP and BACE1, NHE6 overexpression reportedly led to a reduction of Aβ production and conversely shRNA knockdown of NHE6 resulted in an increase in Aβ production (Prasad and Rao, 2015). To investigate if NHE6 deficiency contributes to APP processing by BACE1 in neurons, we used primary neurons derived from APPswe (Tg2576) mice, an Alzheimer’s disease mouse model that overexpresses human APP with the “Swedish” mutation (Hsiao et al., 1996). Neurons were treated with the NHE inhibitor EMD87580 or transduced with lentiviral shRNA directed against NHE6 in the presence or absence of the γ-secretase inhibitor L-685458. βCTF was detected using the monoclonal antibody 6E10 directed against Aβ-residues 1-16 (**Supplemental Figure S2**). Inhibition of γ-secretase in ApoE4-treated APPswe neurons strongly enhanced βCTF accumulation, however, additional treatment with EMD87580 did not alter the amount of βCTF in the cell lysates (**Supplemental Figure S2A**). NHE6 knockdown using lentiviral shRNA also had no effect on the amount of βCTF (**Supplemental Figure S2B**). We conclude that NHE6 inhibition is unlikely to increase Aβ production under near-physiological conditions.

### NHE6 Deficiency Reduces Aβ Plaque Load in Human APP Knockin Mice

To further study the effect of NHE6-deficiency on Aβ pathology *in vivo* we bred humanized APP^NL-F^ mice (Saito et al., 2014) to our germline NHE6 knockout line (NHE6-KO). In these APP^NL-F^ mice, the Aβ-sequence has been completely humanized and the early onset AD Swedish mutation (5’ located mutations encoding K670N and M671L = NL) and the Beyreuther/Iberian mutation (3’ flanking mutation encoding I716F = F) were also introduced, resulting in increased Aβ production, but physiological regulation of APP expression. This allowed us to determine the effect of Aβ overproduction while keeping APP expression under the control of the endogenous promoter. APP^NL-F^;NHE6-KO and control APP^NL-F^ littermates were aged to one year. Perfusion-fixed brains were harvested and analyzed by H&E staining, Thioflavin S staining to visualize plaque load, and Aβ-immunohistochemistry. H&E staining did not reveal any obvious anatomical structural differences between genotypes, but brain size, cortical thickness, hippocampal area, and CA1 thickness were reduced, as described previously (Xu et al., 2017) (**Supplemental Figure S3**). Plaques were more frequent in NHE6 wild type than NHE6-KO mice. To further investigate and quantify plaque load we analyzed the same brains after Thioflavin S staining (**Figure 5A** and **Supplemental Figure S4**). We found an approximate 80% reduction of plaques in NHE6-KO mice when compared to littermate controls (**Figure 5B**). Immunohistochemistry against Aβ showed the same reduction (**Supplemental Figure S5A and S5B**). In addition, we analyzed soluble (TBS fraction) and insoluble (GuHCl and 70% FA fractions) Aβ in cortical brain lysates of 1.5 year old APP^NL-F^;NHE6-KO mice and their control littermates. The amount of insoluble Aβ (GuHCl and 70% FA fractions) was reduced in NHE6-depleted mice by approximately 71%, when compared to their control littermates (**Supplemental Figure S5C-E**). The ∼50% reduction in soluble Aβ was statistically not significant in NHE6-KO lysates (TBS fraction).

**Figure 5.**
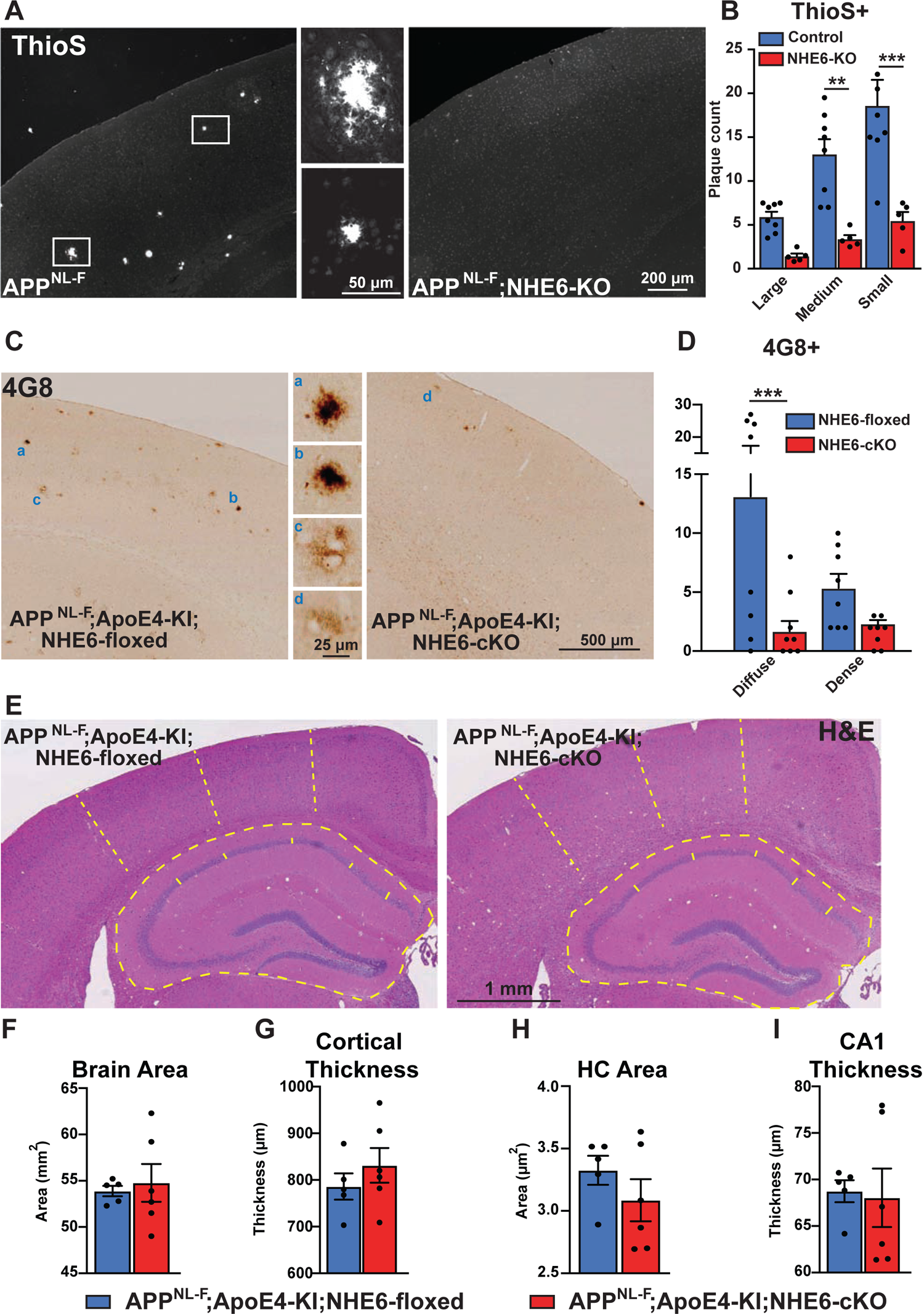
NHE6-Deficiency Decreases Plaque Formation in Both APP^NL-F^ and APP^NL-F^;ApoE4-KI Mice. **(A,B)** NHE6-deficient APP^NL-F^ and control APP^NL-F^ mice were analyzed for plaque deposition at an age of 12 months. Thioflavin S staining was performed to visualize plaques. Plaques were found more frequently in the control APP^NL-F^ mice (left panel in **A**), magnifications of the boxed areas are shown in the two middle panels. The plaque load between NHE6-KO mice and control littermates (all APP^NL-F^) was compared and analyzed. **(B)** In the NHE6-KO littermates the plaque number was reduced, when compared to controls. **(C-D)** NHE6-cKO;APP^NL-F^;ApoE4-KI and NHE6-floxed;APP^NL-F^;ApoE4-KI mice were analyzed for plaque deposition. NHE6 was ablated at two months and brains were analyzed at 13.5-16 months. 4G8-immunolabeling against Aβ was performed to visualize plaques. In APP^NL-F^;ApoE4-KI mice conditional NHE6 knockout caused a reduction in plaque load compared to the NHE6-floxed control littermates (**C**). Magnifications of the boxed areas in C are shown in the middle. **(D)** Plaque load was analyzed and compared between NHE6-cKO mice and floxed control littermates. **(E)** Hematoxylin and eosin staining (H&E) was performed to investigate for gross anatomic abnormalities in the NHE6-cKO;APP^NL-F^;ApoE4-KI and NHE6-floxed;APP^NL-F^;ApoE4-KI mice. **(F-I)** Brain area **(F)**, cortical thickness **(G)**, hippocampal (HC) area **(H)**, and CA1 thickness **(I)** were analyzed. Student *t*-test did not reveal a significant difference. Plaques were differentiated by size or staining density as described in detail in the supplements (Supplement Figure S2). Labeled plaques were analyzed by a blinded observer. All data are expressed as mean ± SEM. **(B)** NHE6-KO n=5, control n=8, NHE6-floxed n=8, NHE6-cKO n=8), in **(F-I)** derived from n=5 (NHE6-floxed) and n=6 (NHE6-cKO) animals. *p < 0.05. **p<0.01, ***p<0.005. Statistical analysis was performed using two-way ANOVA with Sidak’s post-hoc test **(B and D)** and Student *t*-test **(F-I)**.

### NHE6 Deficiency Reduces Plaque Load in APP^NL-F^;ApoE4-KI

To further investigate whether NHE6-deficiency also protects the brain from plaques in the presence of human ApoE4 instead of murine ApoE, we bred NHE6-cKO;ApoE4-KI with APP^NL-F^ mice. At two months of age we induced NHE6-ablation with tamoxifen and aged the mice to 14 - 16 months. NHE6-deficiency on the background of human ApoE4 reduced plaque deposition, as shown by Thioflavin S staining (**Supplemental Figure S5F and S5G**) and 4G8 immunoreactivity **(Figure 5C and 5D)**. APP^NL-F^ mice expressing murine ApoE developed plaques at 12 months (**Figure 5A and 5B**), compared to ApoE4-KI;APP^NL-F^ which showed a similar number of plaques at 15-16 months (**Figures 5C, 5D, 6 and Supplemental Figure S5**). This delay of plaque deposition caused by the presence of human ApoE4 as opposed to murine ApoE is consistent with earlier findings by the Holtzman group (Liao et al., 2015). Importantly, NHE6 ablation induced at two months showed a comparable reduction of plaque load as germline NHE6 depletion. We conclude that plaque deposition is modulated by the presence of NHE6 postnatally and is not affected by NHE6 activity during development.

### NHE6 Deficiency Does Not Affect Cortical Thickness and Hippocampus Size in APP^NL-F^;ApoE4-KI cortices

Xu et al. (2017) reported neuronal loss in the cortex and hippocampus of NHE6-KO mice, which we were able to reproduce in our germline NHE6-KO model (**Supplemental Figure S3**). Neuronal loss can be a trigger for glial activation (Yanuck, 2019). Since both, germline deficiency and adult-onset deficiency of NHE6 causes massive Purkinje cell loss in the cerebellum (**Figure 2C-E**), we next investigated the effect of conditional NHE6-loss on hippocampal and cortical neuronal loss in our APP^NL-F^;ApoE4-KI model (**Figure 5 E-I**). We measured brain size, cortical thickness, hippocampal area, and CA1 thickness. In contrast to germline NHE6-KO mice (**Supplemental Figure S3**) none of the analyzed parameters differed significantly between NHE6-cKO and controls (**Figure 5 E-I**). This indicates that neuronal loss induced by NHE6-deficiency is unlikely to be the trigger for glial activation.

### NHE6 Deficiency Increases Iba1 and GFAP Expression in the Brain

Neuroprotective astrocytes and microglia have been described to reduce Aβ deposition in early stages of AD (Sarlus and Heneka, 2017). It has been reported that NHE6-deficiency leads to increased glial fibrillary acidic protein (GFAP) and ionized calcium-binding adapter molecule (Iba1) immunoreactivity in different brain regions (Xu et al., 2017). To validate if plaque load correlated with Iba1 and/or GFAP immunoreactivity, we performed DAB-immunostaining for both marker proteins. We found that Iba1 and GFAP immunoreactivity is increased in the white matter and to a lesser extent in the cortex of APP^NL-F^;ApoE4-KI;NHE6-cKO and APP^NL-F^;NHE6-KO mice compared to their littermate controls (**Supplemental Figure S6**). There was a non-significant trend towards increased immunoreactivity for both markers in the hippocampus of the NHE6-KO group. Taken together, these data are consistent with the findings of the Morrow group in germline NHE6 KO mice (Xu et al., 2017).

### Microglia and Astrocytes Surround Plaques in Both, NHE6-KO and Control APP^NL-F^ mice

Conditional and germline NHE6-deficient APP^NL-F^ mice have a reduced number of plaques in the brain and an increase of Iba1 and GFAP labeled glia. Others have shown that Aβ plaques levels are reduced with increased plaque-associated microglia detected with a co-stain for Iba1 and Aβ (Parhizkar et al., 2019; Zhong et al., 2019). In order to investigate the contribution of microglia and astrocytes to the observed plaque reduction we analyzed brain slices by costaining for Aβ using the 6E10 antibody (**Figure 7**). Whereas the total amount of plaques labeled by 6E10 was reduced in NHE6-KO as compared to control mice (**Figure 7C**), the amount of microglia and astrocytes surrounding plaques did not differ between genotypes (**Figure 7D and G**). In addition, there was no difference between genotypes in the amount of microglia co-labeled with 6E10 (**Figure 7E**) or the intensity of immunoreactivity for 6E10 within microglia (**Figure 7F**). We hypothesize that the mechanism that leads to reduced plaque load in NHE6-deficient APP^NL-F^ mice may involve an increased catabolic rate (Shi et al., 2021), brought about by the accelerated acidification and vesicular trafficking of early endosomes.

**Figure 6.**
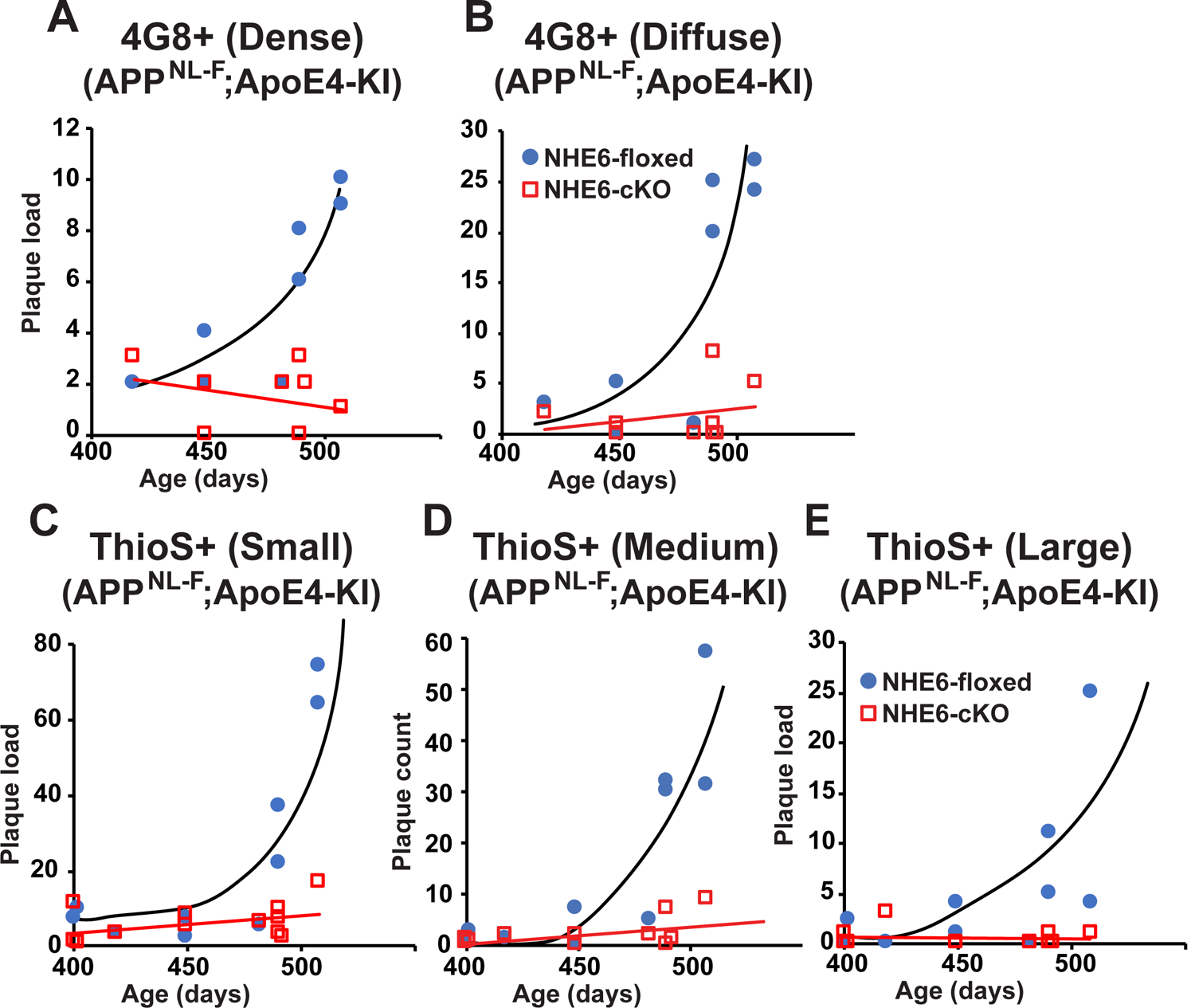
Age Dependent Increase in Plaque Load is abolished in NHE6-cKO;APP^NL-F^;ApoE4-KI Mice. **(A-E)** NHE6-cKO;APP^NL-F^;ApoE4-KI and NHE6-floxed;APP^NL-F^;ApoE4-KI mice were analyzed for plaque deposition. NHE6 was ablated at two months and brains were analyzed at 13.5-16 months. 4G8-immunolabeling against Aβ **(A,B)** and Thioflavin S staining **(C-E)** were performed to visualize plaques (**Figure 5C and Supplement Figure 5F**). Plaque load was analyzed and compared between NHE6-cKO mice and floxed control littermates. Plaques were differentiated by staining intensity **(A, B)** or size **(C-E)** as described in the supplements (**Supplement Figure S4**). In the time range analyzed, plaque load increased by age in control, but not in NHE6-cKO mice. Plaques were analyzed by a blinded observer. Plaque count (NHE6-cKO n=8, NHE6-floxed n=8 for **A** and **B**; NHE6-cKO n=12; NHE6-floxed n=10 in **C-E**) is plotted against age of mice.

**Figure 7.**
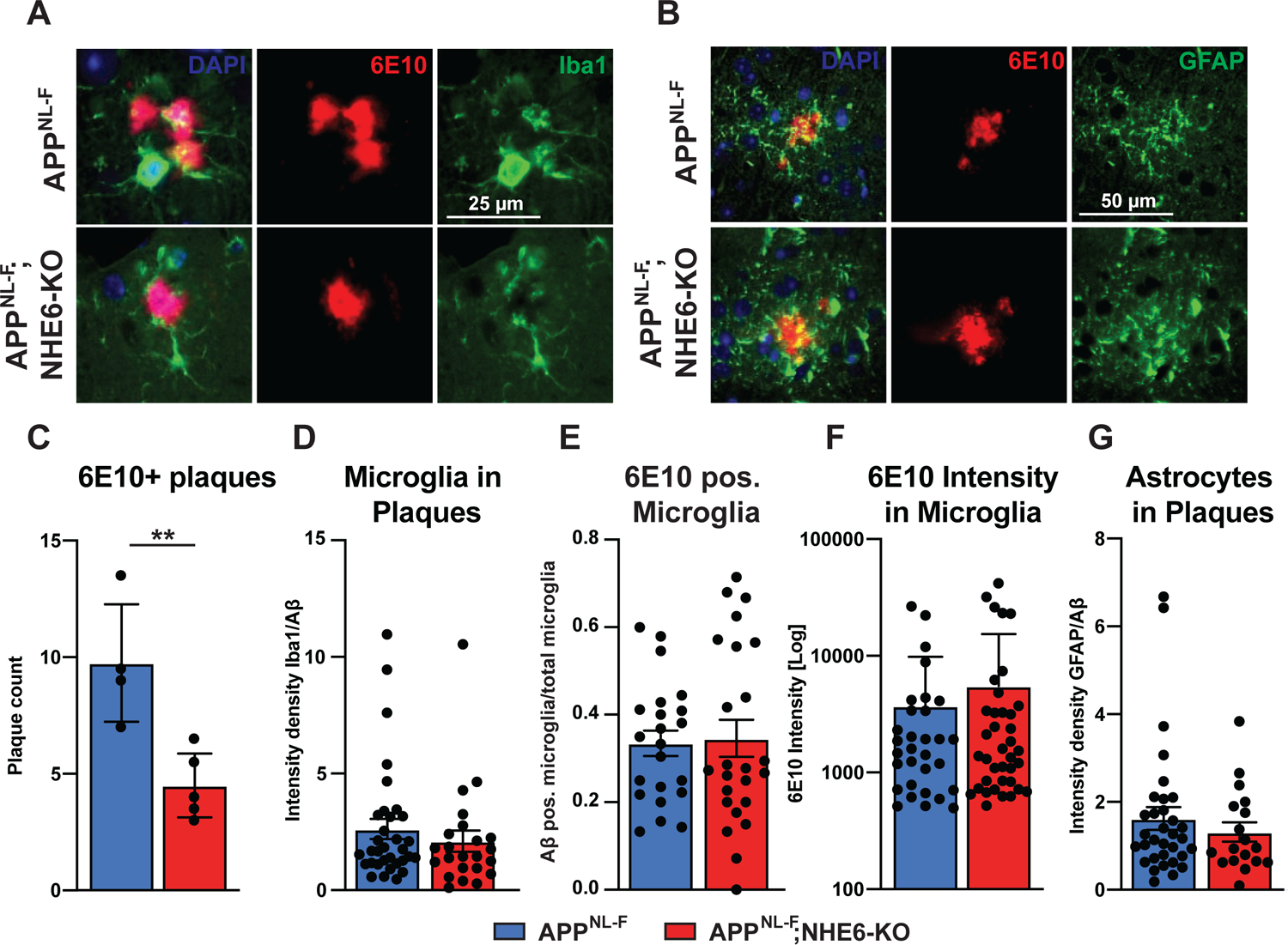
Microglia and Astrocytes Surround Plaques in Both APP^NL-F^ Control and APP^NL-F^;NHE6-KO Brains. **(A-B)** Co-labeling of microglia (Iba1, green, **A**) or astrocytes (GFAP, green, **B**) with Aβ (6E10, red) in brain slices of APP^NL-F^ and APP^NL-F^;NHE6-KO mice. **(C)** Quantification of plaques in control and NHE6-KO brain slices. **(D)** Bar graph showing the intensity density of Iba1/6E10 as quantitative measure of microglia surrounding plaques. **(E)** Statistical analysis of 6E10 positive microglia and **(F)** the intensity of 6E10 signal within microglia. **(G)** Bar graph showing the intensity density of GFAP/Aβ as quantitative measure of astrocytes surrounding plaques. Data were analyzed by a blinded observer. All data are expressed as mean ± SEM. Data were obtained from n=4 (control) and n=5 (NHE6-KO) mice **(A-G)**. **(D)** n=33 (control) and n=23 (NHE6-KO) plaques were analyzed, in **(E)** n=22 (control) and n=24 (NHE6-KO) microscopical pictures were analyzed, in **(F)** n=31 (control) and n=38 (NHE6-KO) 6E10 postive (defined as signal intensity above 500) microglia were analyzed, in **(G)** n=33 (control) and n=18 (NHE6-KO) plaques were analyzed. Student *t*-test revealed a difference in **C** (**p<0.01) and did not reveal significant differences in **D-G**.

## DISCUSSION

The prevalence of Alzheimer’s disease (AD) is increasing with life-expectancy in all human populations. ApoE4 is the most important genetic risk factor. This makes it of paramount importance to understand the underlying mechanisms by which ApoE4 contributes to the pathology of the disease in order to devise effective targeted therapies that can be deployed on a global scale. Only small molecule drug therapies or, alternatively, immunization approaches, can satisfy this requirement. Biologics, including monoclonal antibodies and potential viral gene therapy approaches, are unlikely to be sufficiently scalable. We have previously reported a novel small molecule intervention that has the potential to neutralize ApoE4 risk (Xian et al., 2018) through prevention of the ApoE4-induced endosomal trafficking delay of synaptic receptors by the early endosomal sorting machinery. The mechanistic basis of this conceptually novel intervention is the acidification of the early endosomal compartment through inhibition of NHE6. Remarkably and unexpectedly, loss of NHE6 effectively suppressed amyloid deposition even in the absence of ApoE4, suggesting that hyperacidification of early endosomes occludes the effect of all ApoE forms on amyloid plaque formation. NHE6 suppression or inhibition may thus be a universal approach to prevent amyloid buildup in the brain, irrespective of ApoE genotype. Our previous (Xian et al. 2018) and current studies thus suggest a novel mechanism to prevent ApoE4-risk for Alzheimer’s disease and delay plaque formation. In addition, genome wide association studies (GWAS) in conjunction with studies on cell culture and mouse models of AD show that various AD risk factors enhance endo-lysosomal dysfunction (Knopman et al., 2021; Small et al., 2017; Verheijen and Sleegers, 2018), which potentially could be corrected by NHE6 inhibition.

Upon endocytosis, endosomes undergo gradual acidification controlled by vATPases which actively pump protons into the vesicular lumen, and by the Na^+^/H^+^-exchanger NHE6 which functions as a regulatable proton leak channel. NHE6-depletion acidifies early and recycling endosomes (Lucien et al., 2017; Ohgaki et al., 2010; Ouyang et al., 2013; Xinhan et al., 2011). pH is an important regulator of the endolysosomal sorting machinery in which vesicles undergo multiple rounds of fusion. EEs undergo fusion and fission events in close proximity to the cell membrane. Recycling endosomes originate from EEs while they undergo early-to-late-endosomal maturation. In contrast to late endosomes, recycling endosomes do not undergo further acidification (Jovic et al., 2010; Schmid, 2017). The pH of EEs and recycling endosomes is approximately 6.4-6.5 (Casey et al., 2010). This is normally sufficient to induce ligand receptor dissociation and enable cargo sorting. Our data, however, suggest that ApoE4 dramatically delays this fast recycling step in neurons, where ApoE, Apoer2, and glutamate receptors co-traffic through fast recycling compartments upon Reelin stimulation (Xian et al., 2018). We have proposed that ApoE4 impairs vesicle recycling due to isoelectric precipitation and structural unfolding at the physiological pH of the EE environment. This delays the dissociation of ApoE4 from its receptors, which in turn prolongs the entry of ApoE4 - along with Apoer2 and glutamate receptors in the same vesicle - into the recycling pathway (illustrated in **Figure 1**). We conclude that ApoE4 net-charge affects its endosomal trafficking. This is further supported by recent findings by Arboleda-Velasquez and colleagues (Arboleda-Velasquez et al., 2019), who reported the presence of the “Christchurch” R136S mutation in an E3/E3 PS1 mutation carrier without dementia. By neutralizing the positive charge of Arg136 in ApoE3, the IEP of this ApoE3 isoform is predicted to match that of ApoE2, which is protective against AD (Corder et al., 1994). ApoE2 homozygous carriers have an exceptionally low likelihood of developing AD (Reiman et al., 2020). Moreover, the Christchurch mutation is located within the heparin binding domain of ApoE, which reduces its affinity for cell surface heparan sulfate proteoglycans. That in turn would result in decreased uptake and thus depletion of ApoE in EEs. The net effect would be unimpeded trafficking of EE vesicles through the fast recycling compartment.

In dendritic spines, NHE6 co-localizes with markers of early and recycling endosomes and with the glutamate receptor subunit GluA1. In the hippocampus, NHE6 is highly expressed in the pyramidal cells of the CA and the granule cells of the dentate gyrus (Stromme et al., 2011). Apoer2 is present at the postsynaptic density of CA1 neurons (Beffert et al., 2005). During LTP induction, translocation of NHE6-containing vesicles to dendritic spine heads is enhanced (Deane et al., 2013) and glutamate receptors are recruited to the synaptic surface through fast recycling (Fernandez-Monreal et al., 2016). Our findings are consistent with a model where NHE6 serves as a pH-regulator of Reelin-controlled fast recycling endosomes containing Apoer2 and glutamate receptors. This mechanism possibly translates to other cell types and other ApoE receptors.

We previously showed that ApoE4 impairs Reelin-mediated receptor recruitment to the neuronal surface and this can be reversed by functionally disabling NHE6 in primary neurons, which results in the increased acidification of EEs to a level sufficiently different from the IEP of ApoE4, which then allows its efficient dissociation from its receptors. Conditional NHE6 deletion accordingly alleviates the ApoE4-mediated resistance to Reelin-enhanced synaptic plasticity in hippocampal field recordings.

Amyloid-β (Aβ) and tau, forming amyloid plaques and neurofibrillary tangles, are the defining features of AD pathology. As of today, it remains controversial how ApoE-isoforms interfere with Aβ and tau pathology. ApoE, which is primarily expressed by astrocytes, is the major lipid transporter in the brain and in an isoform-dependent manner affects inflammatory, endolysosomal and lipid-metabolic pathways (Gao et al., 2018; Minett et al., 2016; Van Acker et al., 2019; Xian et al., 2018). Most risk factors identified by genome-wide association studies, including but not limited to APOE, ABCA7, CLU, BIN1, TREM2, SORL1, PICALM, CR1 are members of one or more of these pathways (Kunkle et al., 2019). In recent years, endosomal dysfunction has increasingly gained acceptance as a causal mechanism for late-onset AD. Our findings now provide a mechanistic explanation how ApoE4 impairs endolysosomal trafficking and recycling, by interfering with vesicular sorting and maturation at a crucial bottleneck juncture of the endosomal trafficking machinery. This has far-reaching consequences for neuronal function, synaptic plasticity, and tau phosphorylation (Brich et al., 2003; Cataldo et al., 2000; Chen et al., 2005; Chen et al., 2010; Nuriel et al., 2017; Pensalfini et al., 2020). More specifically, ApoE4 causes abnormalities of Rab5-positive endosomes (Nuriel et al., 2017). Intriguingly, over-activation of the small guanosine triphosphatase (GTPase) Rab5, recapitulates neurodegenerative features of AD (Pensalfini et al., 2020).

ApoE4 alters APP processing and Aβ-degradation, (reviewed in (Benilova et al., 2012; Haass et al., 2012; Huynh et al., 2017; Lane-Donovan and Herz, 2017; Pohlkamp et al., 2017; Yamazaki et al., 2019)) and the ability of Reelin to protect the synapse from Aβ toxicity is impaired by ApoE4 (Durakoglugil et al., 2009). Aβ-oligomerization followed by plaque formation is one hallmark of AD. NHE6 controls endosomal pH, which can affect BACE1 activity, one of the two enzymes required to process APP to release the Aβ-peptide. Prasad and Rao (2018) reported that overexpression of NHE6 in astrocytes, rather than its inhibition or knockdown, reduced Aβ generation, which conflicts with our findings in primary cortical neurons. Although the cause of this discrepancy remains currently unresolved, it is possible that it is the result of the two fundamentally different experimental systems that were used in the respective studies, i.e. overexpression in tissue culture on one hand and genetic manipulation of live mice on the other. Using a humanized APP^NL-F^ knockin mouse model we show that NHE6-deficiency in one-year old animals reduces plaque deposition by approximately 80%. In APP^NL-F^ mice plaques can be identified as early as 9 months of age. Whereas plaque deposition only increases by less than twofold between 9 and 12 months of age, it increases tenfold between 12 and 18 months (Saito et al., 2014). Importantly, the reduction in plaque deposition by NHE6-deficiency persists from early (12 months) to later stages (18 months) of AD, as NHE6-deficient APP^NL-F^ animals aged 18 months had a reduction in insoluble Aβ by approximately 71%. NHE6 depletion in APP^NL-F^;ApoE4-KI mice showed a comparable reduction in plaque load (**Figure 5 and Supplemental Figure S5**). Our data are consistent with previous findings by the Holtzman group (Fagan et al., 2002) that showed that mouse ApoE promotes plaque deposition more potently than human ApoE4.

Prevention of plaque formation in our NHE6 deficient model was likely caused by increased microglial activation and plaque phagocytosis (**Figure 5, 6, 7, Supplemental Figure S5, S6 and S7**). In the brains of AD patients and APP overexpressing mice, plaques are surrounded by reactive microglia and astrocytes (Meyer-Luehmann et al., 2008; Serrano-Pozo et al., 2013), but the pathological significance of this is incompletely understood. Beneficial or detrimental roles of reactive microglia and astrocytes in the degradation of Aβ have been reported, depending on the activation state of these cells (Meyer-Luehmann et al., 2008; Ziegler-Waldkirch and Meyer-Luehmann, 2018). We observed an increase in reactive microglia and astrocytes resulting from NHE6-depletion, which correlated with reduced plaque deposition in APP^NL-F^ mice, irrespective of the presence of either murine ApoE or human ApoE4. As murine ApoE exacerbates plaque deposition even more than ApoE4, the comparable plaque reduction in NHE6 deficient mice with murine ApoE or ApoE4-KI might be the result of a maximally accelerated early endosomal maturation and cargo transport in the absence of NHE6. When compared to control APP^NL-F^ mice, NHE6-deficient APP^NL-F^ mice show an increase in Iba1 (microglia) and GFAP (astrocytes) immunoreactivity, but reduced Aβ immunoreactivity. Surprisingly, the intensity of GFAP and Iba1 in plaque areas was comparable between the groups. Moreover, even though Aβ is reduced and Iba1 is increased in the NHE6-KO, the proportion of microglial structures containing Aβ (6E10 antibody) was comparable between NHE6 deficient and control APP^NL-F^ mice, as was the intensity signal for 6E10. This suggests that microglia in the NHE6-KO may be more efficient in taking up and degrading Aβ. Whether the reduction in plaques is due to the presence of an increased number of microglia and astrocytes that actively phagocytose nascent plaques, or whether endosomal acidification in microglia and astrocytes improves their ability to degrade or export Aβ from the brain remains to be determined. It is also possible that NHE6-deficiency alters the efficiency of astrocytes to lipidate ApoE. In a mouse model, improved ApoE lipidation by the overexpression of ATP-binding cassette transporter family member A1 (ABCA1) decreased plaque deposition (Wahrle et al., 2005; Wahrle et al., 2008; Wahrle et al., 2004). During HDL assembly, ABCA1 shuttles between EE and the plasma membrane, a process also referred to as retroendocytosis (Ouimet et al., 2019). Moreover, membrane trafficking of ABCA1 is altered by ApoE in an isoform dependent fashion (Rawat et al., 2019). The APP^NL-F^ mouse model used in our study develops plaques at 12 months (Saito et al., 2014) in the presence of murine ApoE. However, onset of plaque deposition in human ApoE4-KI mice was delayed by approximately three months. The effect of germline NHE6 deficiency and conditional NHE6-deficiency induced at two months had a comparable effect on plaque reduction in APP^NL-F^ and APP^NL-F^;ApoE4-KI mice.

NHE6-KO and NHE6-cKO both show progressive Purkinje cell loss in the cerebellum, indicating that NHE6 requirement is cell-autonomous and not developmentally determined. NHE6-KO and NHE6-cKO show a comparable increase in immunoreactivity against Iba1 and GFAP. Increased glia reactivity can be a direct cell-autonomous effect of NHE6-loss or an indirect effect caused by NHE6-deficiency related neuronal death. As germline NHE6-KO mice present with cortical and hippocampal neuronal loss (Xu et al., 2017) (**Supplemental Figure S3**) it is possible that it is this neuronal cell death that gives rise to glia activation. However, the tamoxifen induced NHE6-cKO mice do not present with cortical or hippocampal neuronal loss (**Figure 5**), yet have comparable immunoreactivity for markers of glial activation. Moreover, this further indicates that reduced plaque load is also not an effect of neuronal loss. Therefore, our two mouse models together suggest that the observed increased glial activation is not caused on neuronal cell loss but rather is likely a direct cell-autonomous effect of NHE6 loss of function. It thus remains to be determined whether endosomal acidification in NHE6 deficient microglia alone is sufficient to induce Aβ degradation and plaque reduction. It would also be intriguing to test whether NHE6-deficiency increases cognitive performance in AD mouse models. In the current study we used APP^NL-F^ and APP^NL-F^;ApoE4-KI mice as the most physiological currently available mouse models of AD that do not rely on excessive amyloid overproduction. These mice, however, do not show cognitive impairments in spatial learning tests like Morris water maze (Saito et al., 2014) (own unpublished observations). Other neurobehavioral phenotypes have been described for NHE6-KO mice, which also recapitulate symptoms in Christianson syndrome patients, for example hyposensitivity to pain (Petitjean et al., 2020). Future studies on our novel tamoxifen inducible NHE6-cKO line will help to understand whether these symptoms are based on neurodevelopmental deficits caused by germline NHE6 deficiency or whether they can be reproduced by induced loss of NHE6 postnatally.

In conclusion, we have shown that both, the endosomal trafficking defect induced by ApoE4 in neurons as well as increased plaque deposition irrespective of ApoE genotype can be corrected by inhibition or genetic deletion of NHE6, a key regulator of early endosomal pH. Accelerated acidification of early endosomes abolishes the ApoE4-induced Reelin resistance and restores normal synaptic plasticity in ApoE4 targeted replacement mice. The first FDA approved drug for AD treatment in 18 years is aducanumab (Sevigny et al., 2016), an antibody directed against Aβ, which clears amyloid from the brain. However, amyloid removal in individuals already afflicted with AD provides at best marginal benefits at this stage. Moreover, in excess of 1 billion people world-wide are ApoE4 carriers, making early treatment with a complex biologic such as aducanumab impractical on the global scale. Here we have presented a potential alternative approach which should be adaptable to large-scale prevention treatment using blood-brain-barrier penetrant NHE6-specific inhibitors. Taken together, our combined data suggest that endosomal acidification has considerable potential as a novel therapeutic approach for AD prevention and possibly also for the prevention of disease progression.

## Acknowledgements

This work was supported by NIH grants R37 HL063762, R01 NS093382, R01 NS108115, and RF1 AG053391 to JH and 1F31 AG067708-01 to CHW as well as funding from the Darrell K. Royal Research Fund to MD. While this work was ongoing, JH was further supported by the Bright Focus Foundation (A20135245) & (A2016396S); Harrington Discovery Institute; & Circle of Friends Pilot Synergy Grant; and the Blue Field Project to Cure FTD. We are indebted Rebekah Hewitt, Barsha Subbha, Huichuan Reyna, Issac Rocha, Tamara Terrones, Emily Boyle, Alisa Gilloon, Travis Wolff, and Eric Hall for their excellent technical assistance. We thank Dr Yuan Yang for creating the NHE6-FLAG plasmid and the UTSW Whole Brain Microscopy Facility (WBMF) in the Department of Neurology and Neurotherapeutics for assistance with slide scanning. The WBMF is supported by the Texas Institute for Brain Injury and Repair (TIBIR). John Shelton and the UT Southwestern’s Histopathology Core provided help with paraffin sectioning as well as H&E and Thioflavin S staining. We thank Wolfgang Scholz for providing EMD87580.

## Author Contribution

Theresa Pohlkamp, designed and performed research, designed and created the unpublished NHE6-KO/cKO mouse lines, analyzed data, and wrote the paper. Xunde Xian, designed and performed research, analyzed data, reviewed and edited the paper. Connie H Wong, designed and performed research, analyzed data, illustrated figures, reviewed and edited the paper. Murat S Durakoglugil, designed and performed research, analyzed data, reviewed and edited the paper. Takaomi Saido provided the humanized APP^NL-F^ mice. Jade Connor, performed research. Bret M Evers, performed research. Charles L White, performed research. Robert E Hammer, performed mouse manipulations to create the new NHE6-floxed mouse line. Joachim Herz, conceptualization, research design, resources, formal analysis, supervision, funding acquisition, validation, investigation, methodology, writing and editing the paper.

---

**Supplemental Figure S1.**
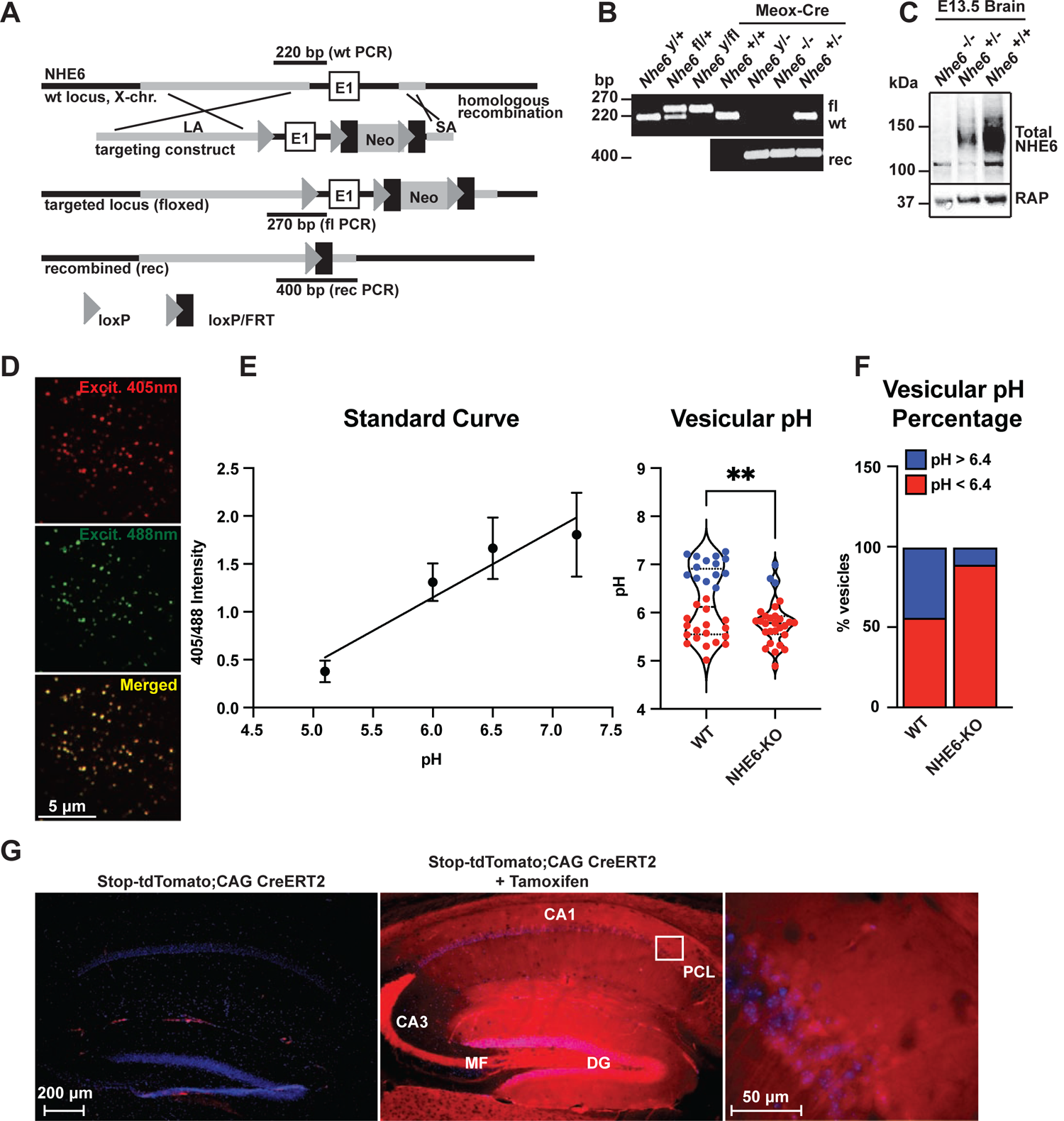
Generation of NHE6-Floxed and NHE6-KO Mice. **(A)** Gene targeting strategy. LoxP sites were introduced to flank the first exon (E1) of NHE6 (located on the X-Chromosome) by gene targeting in embryonic stem cells. The targeting construct contained a long arm of homology (LA, grey) upstream of the first loxP site and the first exon. A loxP/FRT-flanked neomycin resistance cassette was cloned downstream of the first exon, followed by a short arm of homology (SA, grey). The targeted locus is shown below. Targeted stem cells were used to generate chimeric NHE6-floxed mice. Germline NHE6 knockout mice (NHE6-/-(female), NHE6y/-(male); rec indicates recombined allele) were generated by breeding the NHE6-floxed line with Meox-Cre mice. **(B)** Genotyping of wildtype (wt, +), floxed (fl), and recombined (rec, -) NHE6 alleles. The PCR amplified regions are indicated in panel A. The wildtype and floxed allele PCR products differ by 50 bp (270 for floxed, 220 for wildtype). **(C)** Western blot showing brain lysates (left) of different NHE6 genotypes after Meox-Cre induced germline recombination. **(D)** Mouse embryonic fibroblasts from NHE6-KO and control littermate were infected with Vamp3-pHluorin2 and excited at 408 and 488 nm with emission measured at 510 nm. **(E)** vesicular pH measured using a standard curve was significantly decreased in NHE6-KO fibroblasts. **(F)** The percent of vesicles with pH >6.4 is significantly decreased in NHE6-KO fibroblasts. **(G)** CAG-CreERT2 activity after tamoxifen application in a reporter mouse line expressing tdTomato. CreERT2 recombination activity without (left panel) or with (middle panel) tamoxifen application in the CAG-CreERT2-line bred with Rosa26-stop-tdTomato line. After tamoxifen induction CreERT2 activity led to a robust tdTomato signal in the hippocampus (middle panel). Pyramidal neurons in the CA1 pyramidal cell layer (PCL) (middle panel) are shown magnified in the right panel.

**Supplemental Figure S2.**
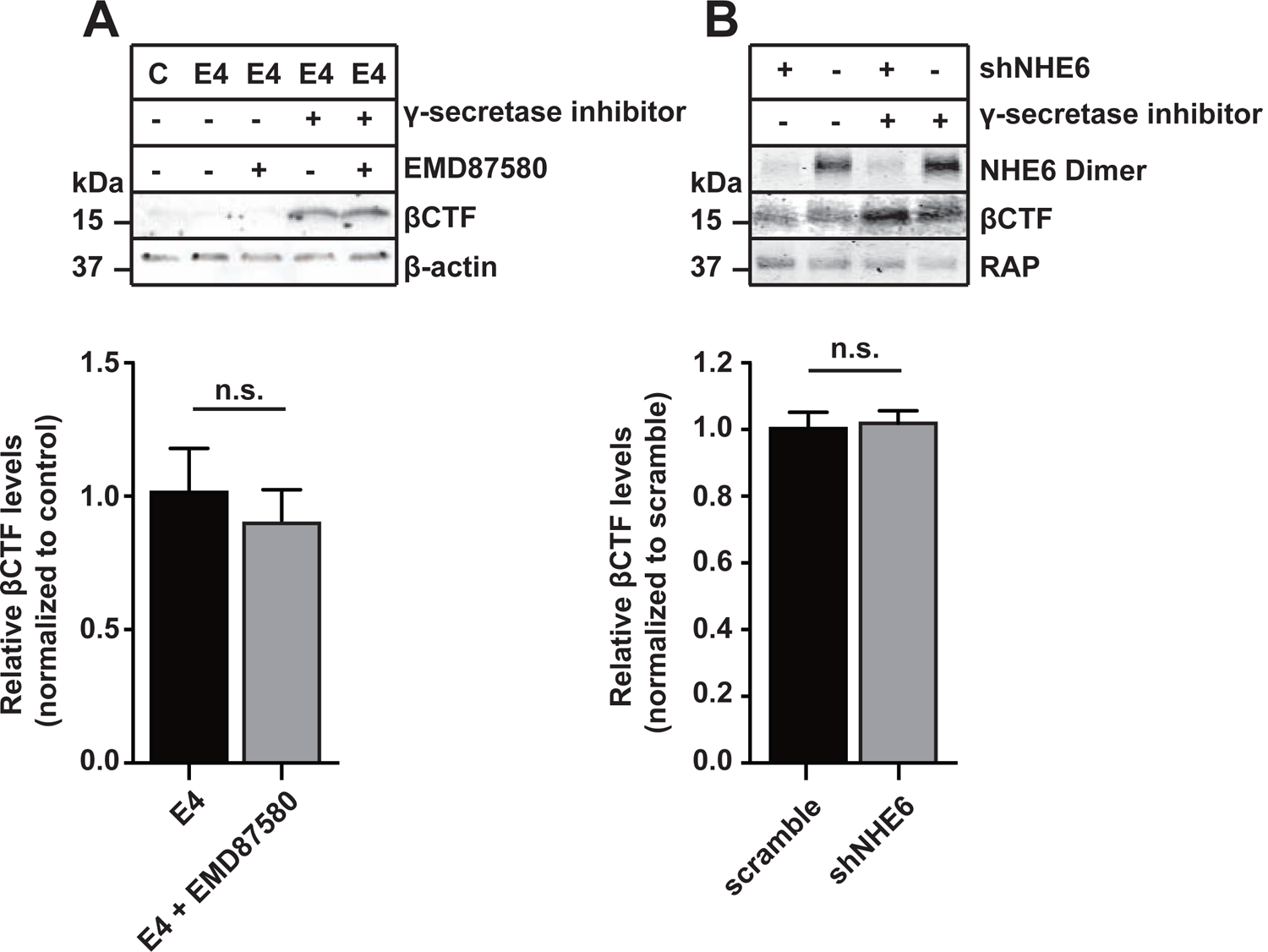
NHE Inhibition or NHE6 Knockdown Does Not Alter BACE1 Activity in Primary Neurons. **(A,B)** Pan-NHE inhibition by EMD87580 or lentiviral knockdown of NHE6 did not alter BACE1 activity in primary neurons of APPswe mice (Tg2576). **(A)** DIV10 primary neurons were treated with γ-secretase inhibitor L-685458, EMD87580, and/or ApoE4 (as indicated) and harvested for immunoblotting against Aβ-containing C-terminal fragment of APP (βCTF). β-actin was blotted as loading control. Bar graph shows the statistics of n = 3 experiments. **(B)** Primary neurons of APPswe mice were infected with lentivirus for shRNA expression directed against NHE6 (shNHE6) or a scramble control sequence (-) at DIV7. At DIV13 neurons were treated with L-685458 over night and harvested for immunoblotting against NHE6 and βCTF on DIV14. RAP was blotted as loading control. Bar graph shows the statistics of n = 6 experiments. All data are expressed as mean ± SEM. Statistical analysis was performed using Student *t*-test. n.s. = not significant.

**Supplemental Figure S3.**
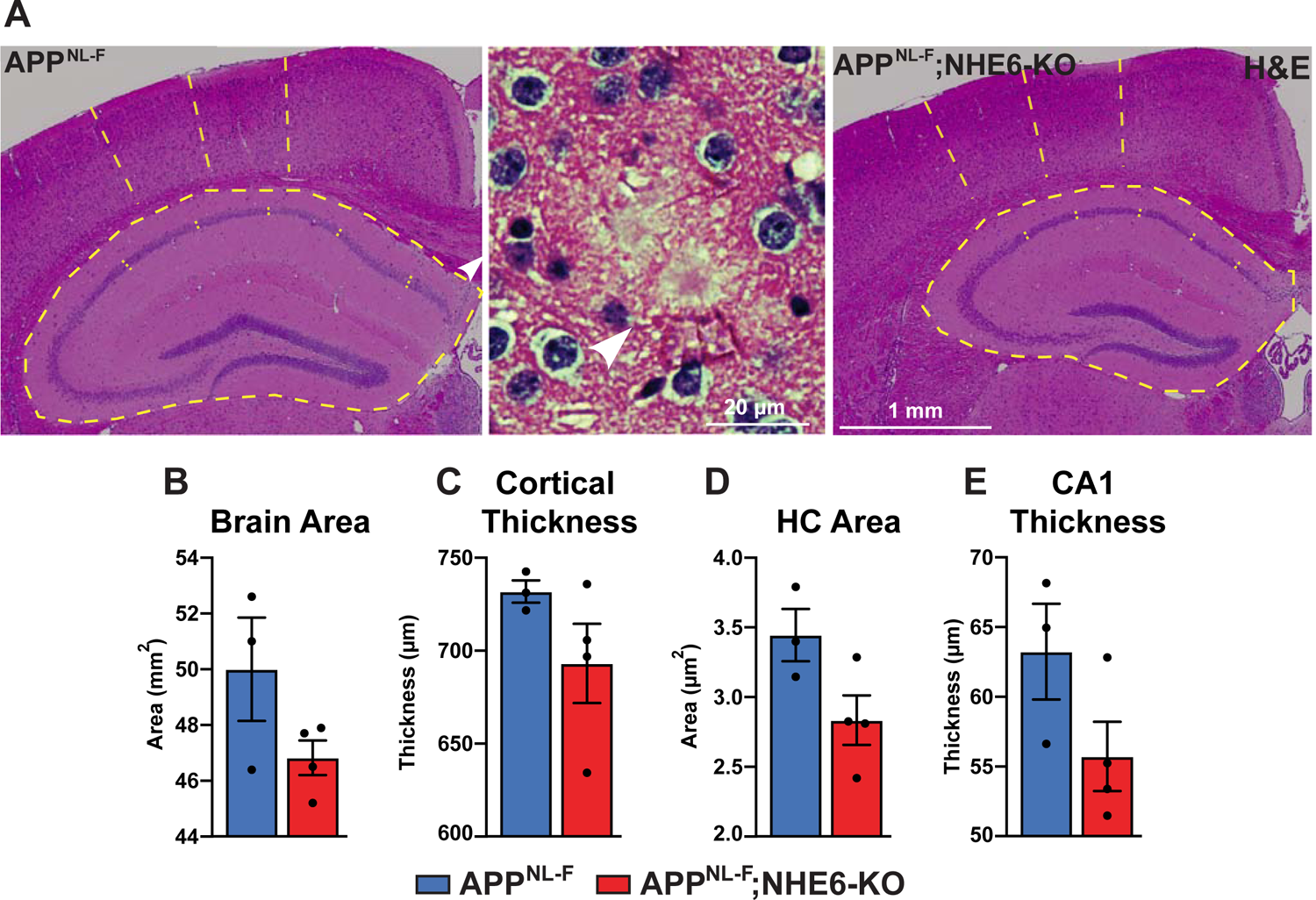
Gross Anatomical Brain Structure in NHE6-KO Mice. **(A)** Hematoxylin and eosin staining (H&E) was performed to investigate for gross anatomic abnormalities in the NHE6-KO;APP^NL-F^ and APP^NL-F^ mice. Structures representing plaques were found in the APP^NL-F^ control groups (magnified example is shown in the middle panel). **(B-E)** Brain area **(B)**, cortical thickness **(C)**, hippocampal (HC) area **(D)**, and CA1 thickness **(E)** were analyzed. All data are expressed as mean ± SEM. Student *t*-test did not reveal significant differences, n=3 (NHE6-KO) and n=4 (control).

**Supplemental Figure S4.**
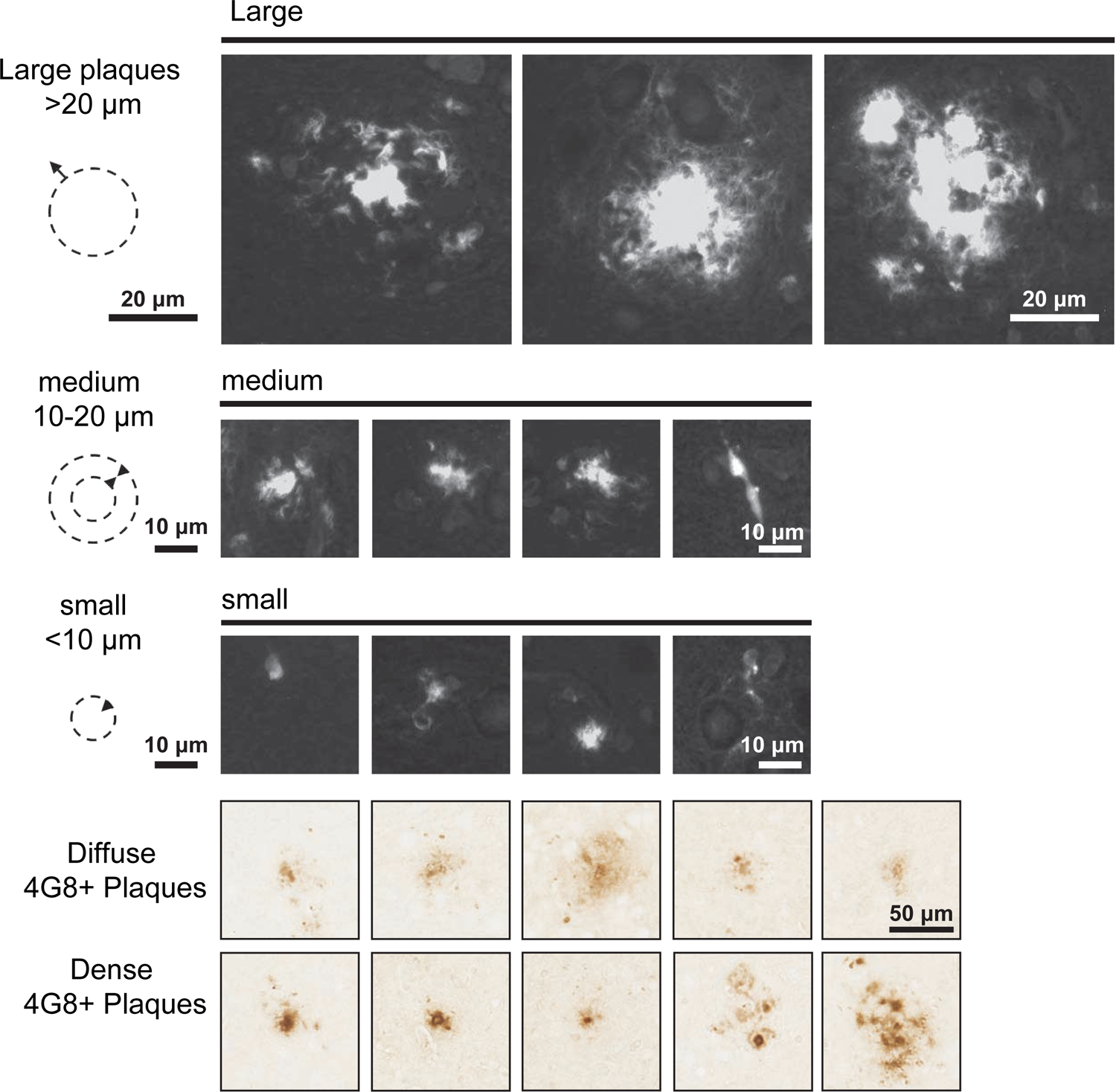
Example of Thioflavin S Stained Plaques for Quantification. Different types of Thioflavin S stained plaques and 4G8-immunoreactive accumulations in APP^NL-F^ brains are shown. Different sizes of plaques were grouped together for quantification Thioflavin S labeled plaques (**Figure 6D**). Plaques bigger in diameter than 20µm with a dense core were defined as big. Medium sized plaques had a diameter between 10-20µm with a dense core. Small plaques were smaller than 10µm and often represented individual cells. 4G8-labeled plaques were differentiated by diffuse or dense appearance as depicted in the examples.

**Supplemental Figure S5.**
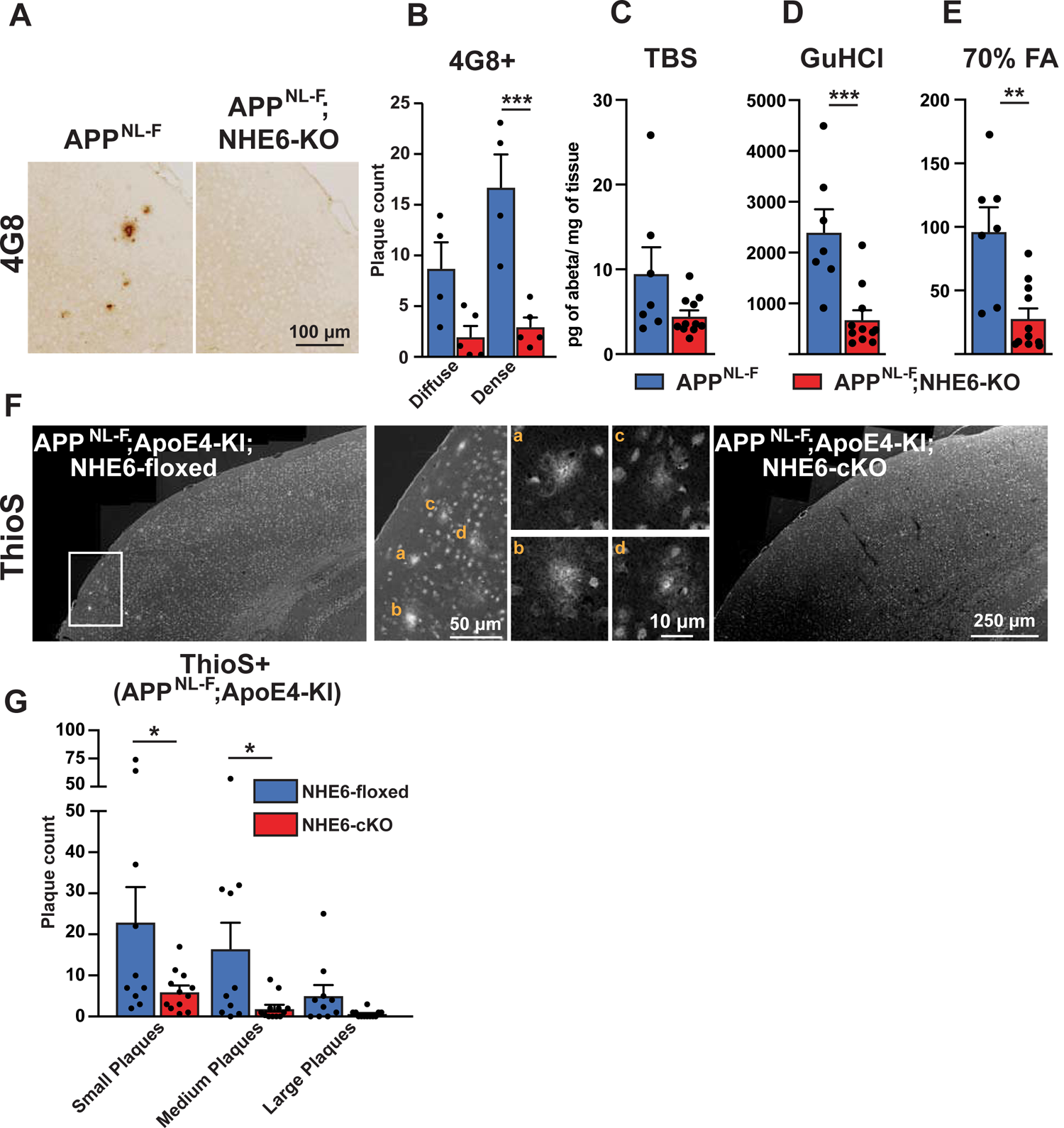
Additional Material to Figure 5 and 6. NHE6-Deficiency Decreases Plaque Formation in Both APP^NL-F^ and APP^NL-F^;ApoE4-KI Mice. **(A-B)** NHE6-deficient APP^NL-F^ and control APP^NL-F^ mice were analyzed for plaque deposition at an age of 12 months. 4G8-immunolabeling against Aβ **(A)** visualized more plaques in the control APP^NL-F^ mice. The plaque load between NHE6-KO mice and control littermates (all APP^NL-F^) was compared and analyzed **(B).** In the NHE6-KO littermates the plaque number was reduced, when compared to controls. **(C-E)** Soluble (TBS) and insoluble (GuHCl and 70% FA) Aβ fractions of cortical lysates were analyzed by commercial ELISA. 1.5-year-old NHE6-KO mice showed less insoluble Aβ than their control littermates (all APP^NL-F^). **(F-G)** NHE6-cKO;APP^NL-F^;ApoE4-KI and NHE6-floxed;APP^NL-F^;ApoE4-KI mice were analyzed for plaque deposition. NHE6 was ablated at two months and brains were analyzed at 13.5-16 months. Thioflavin S staining was performed to visualize plaques. With APP^NL-F^;ApoE4-KI background the NHE6-cKO mice had a reduced plaque load compared to the NHE6-floxed control mice (left panel in **F**). Magnifications of the boxed areas in left panel are shown in the middle. **(G)** Plaque load was analyzed and compared between NHE6-cKO mice and floxed control littermates. Plaques were differentiated by size or staining density as described in detail in the supplements (**Supplement Figure S2**). Labeled plaques were analyzed by a blinded observer **(B, G)**. All data (immunohistochemistry: NHE6-KO n=5, control n=4, NHE6-floxed n=10, NHE6-cKO n=12; biochemistry: NHE6-KO n=11, control n=7) are expressed as mean ± SEM. *p < 0.05. **p<0.01, ***p<0.005. Statistical analysis was performed using two-way ANOVA with Sidak’s post-hoc test.

**Supplemental Figure S6.**
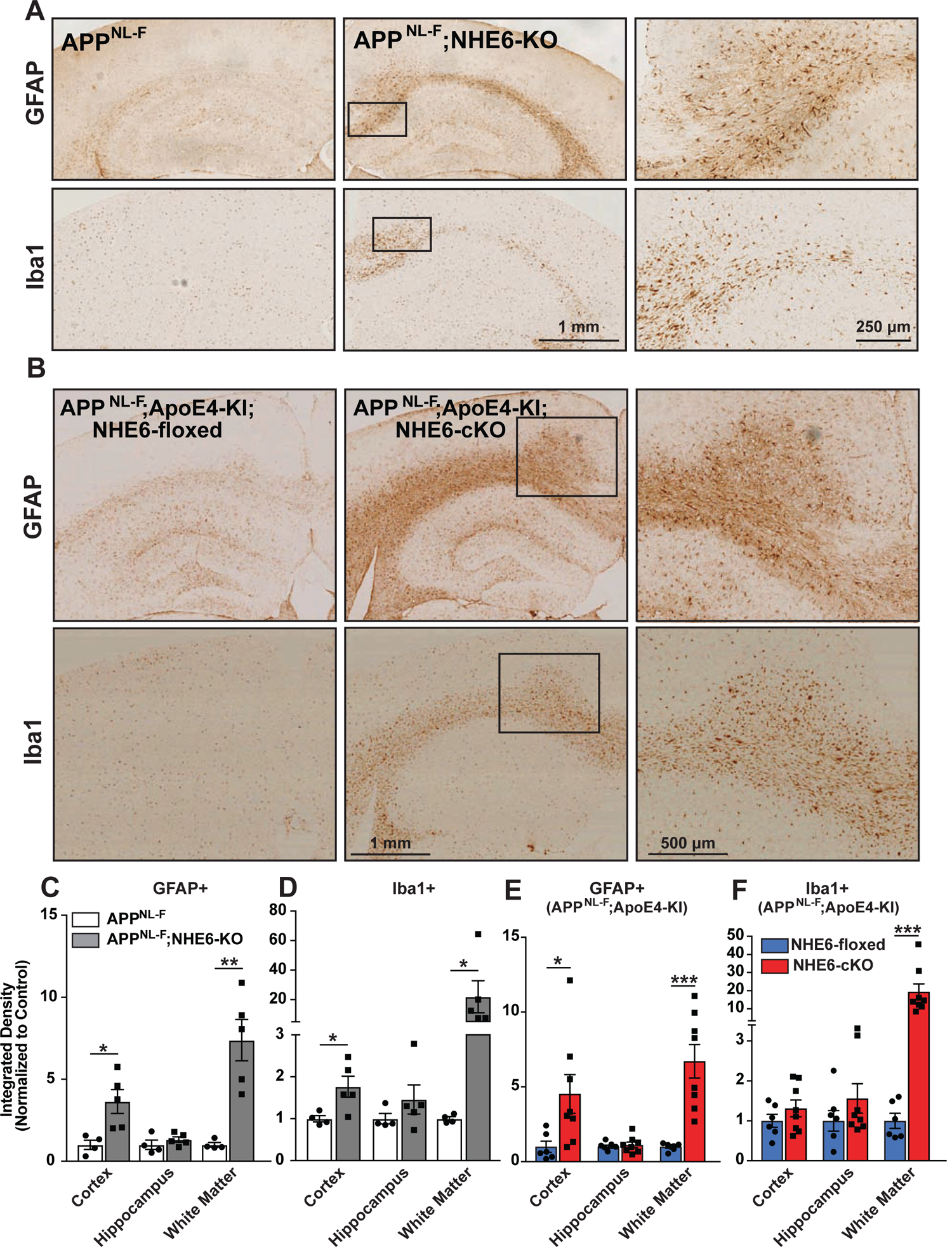
NHE6-Deficiency Causes an Increase in Iba1 and GFAP Immunoreactivity in Both APP^NL-F^ and APP^NL-F^;ApoE4-KI mice. **(A)** Immunohistochemistry against glial fibrillary acidic protein (GFAP) and ionized calcium-binding adapter molecule (Iba1) was performed on brain slices obtained from one-year-old APP^NL-F^ mice deficient for NHE6 (APP^NL-F^;NHE6-KO) and control littermates (APP^NL-F^). GFAP (upper panels) and Iba1 (lower panels) immunoreactivity was increased in NHE6-deficient APP^NL-F^ mice when compared to NHE6 expressing controls. **(B)** NHE6-cKO;APP^NL-F^;ApoE4-KI and NHE6-floxed;APP^NL-F^;ApoE4-KI mice were analyzed for immunoreactivity GFAP and Iba1. Mice were injected with tamoxifen at two months and brain slices obtained from 13.5-16 months old mice. GFAP (upper panels) and Iba1 (lower panels) immunoreactivity was increased in NHE6-deficient APP^NL-F^;ApoE4-KI mice when compared to NHE6 expressing controls. **(C-F)** Intensity of the staining in various areas was compared for GFAP **(C)** and Iba1 **(D)** in APP^NL-F^;NHE6-KO and APP^NL-F^ mice. Intensity of the staining for GFAP **(E)** and Iba1 **(F)** in NHE6-cKO;APP^NL-F^;ApoE4-KI and NHE6-floxed;APP^NL-F^;ApoE4-KI mice. Analysis was performed by a blinded observer. ‘White matter’ comprises corpus callosum, cingulum and external capsule. All data (NHE6-KO n=5; control n=4; NHE6-cKO n=6; NHE6-floxed n=8) are expressed as mean ± SEM. *p < 0.05. **p<0.01, ***p<0.005. Statistical analysis was performed using Student *t*-test.

**Supplemental Figure S7.**
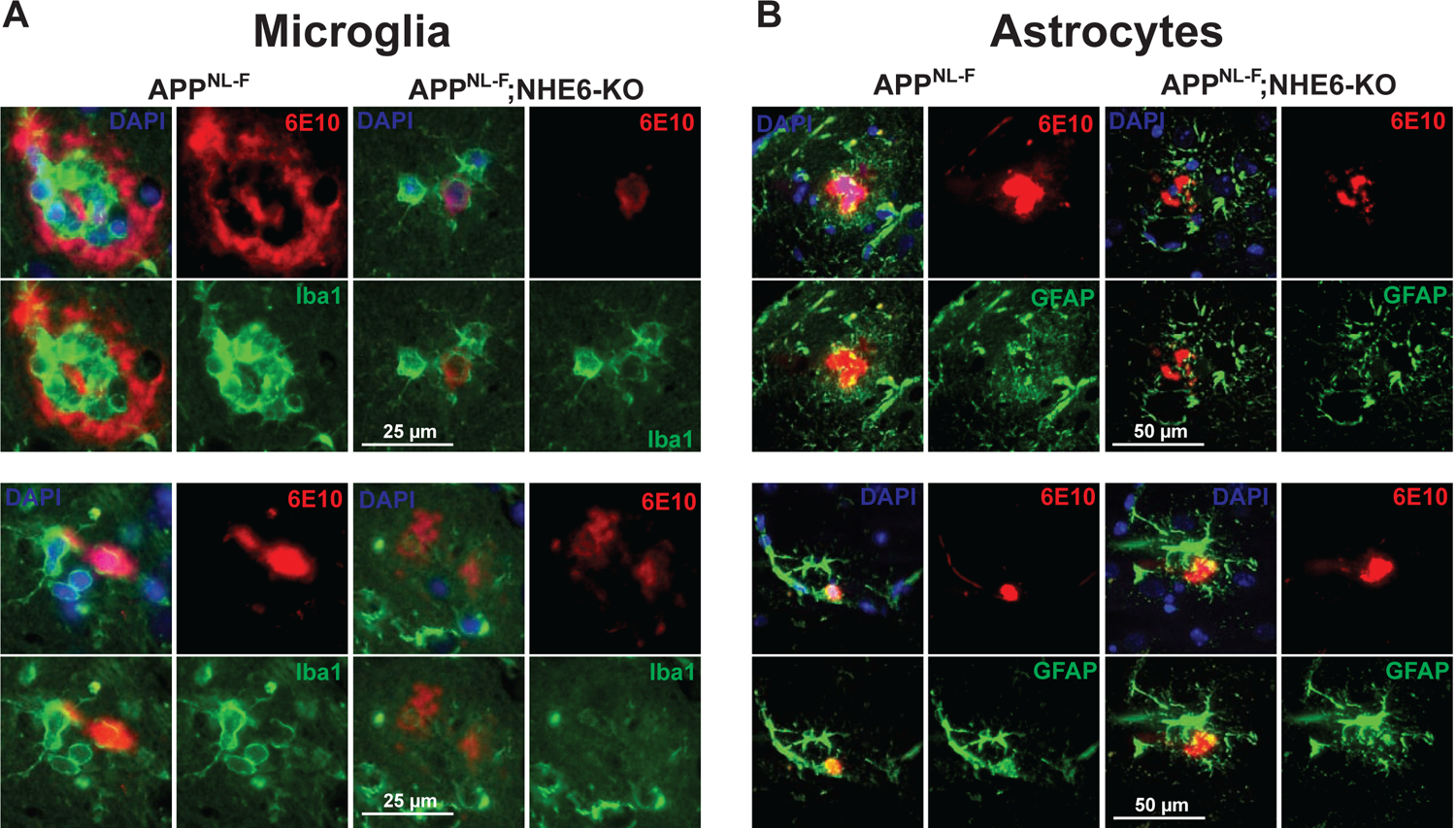
Examples of Plaques Surrounded by Microglia and Astrocytes. Co-labeling of microglia (Iba1, green) and Aβ (6E10, red) in brain slices of APP^NL-F^ and APP^NL-F^;NHE6-KO mice. **(B)** Co-labeling of astrocytes (GFAP, green) and Aβ (6E10, red) in brain slices of APP^NL-F^ and APP^NL-F^;NHE6-KO mice. (Supplementing Figure 7 of the main manuscript).

